# Ultrafast (>400 Hz) network oscillations induced in thalamorecipient cortical layers by optogenetic activation of thalamocortical axons

**DOI:** 10.1101/2022.09.02.506344

**Authors:** Hang Hu, Rachel E. Hostetler, Ariel Agmon

**Affiliations:** Dept. of Neuroscience, West Virginia University School of Medicine, WV Rockefeller Neuroscience Institute, Morgantown, WV 26506, USA

## Abstract

Oscillations of extracellular voltage, reflecting synchronous rhythmic activity in large populations of neurons, are a ubiquitous feature in the mammalian brain and are thought to subserve critical, if not fully understood cognitive functions. Oscillations at different frequency bands are hallmarks of specific brain or behavioral states. At the higher end of the scale, ultrafast (400-600 Hz) oscillations in the somatosensory cortex, in response to peripheral stimulation, were observed in human and a handful of animal studies; however, their synaptic basis and functional significance remain largely unexplored. Here we report that brief optogenetic activation of thalamocortical axons ex-vivo elicited precisely reproducible, ~410 Hz local field potential wavelets (“ripplets”) in middle layers of mouse somatosensory (barrel) cortex. Fast-spiking (FS) inhibitory interneurons were exquisitely synchronized with each other and fired spike bursts in anti-phase with ripplets, while excitatory neurons fired only 1-2 spikes per stimulus. Both subtypes received shared excitatory inputs at ripplet frequency, and bursts in layer 5 FS cells required intact connection with layer 4, suggesting that layer 4 excitatory cells were driving FS bursts in both layers. Ripplets may be a ubiquitous cortical response to exceptionally salient sensory stimuli, and could provide increased bandwidth for encoding and transmitting sensory information. Lastly, optogenetically-induced ripplets are a uniquely accessible model system for studying synaptic mechanisms of fast and ultrafast oscillations.

## INTRODUCTION

A ubiquitous feature of neuronal networks, in the brains of multiple mammalian species throughout the evolutionary tree, is their propensity to engage in large-scale, synchronous and rhythmic patterns of electrical activity, reflected as oscillations in the local field potential (LFP) (Buzsaki et al., 2013). LFP oscillations occur at frequencies spanning 4 or more orders of magnitude, with oscillations at different frequency bands often nested together and co-modulating each other (Steriade, 2006). Specific oscillatory patterns are hallmarks of specific brain or behavioral states; for example, theta rhythms (4-9 Hz) are typical of exploratory behavior, sleep spindles (~10 Hz) occur during certain sleep stages, and gamma oscillations (30-90 Hz) are the hallmark of cognitive activity and sensory processing (Lopes da Silva, 2013; Singer, 2018). Even faster, transient 150-200 Hz “ripples” in the pyramidal cell layer of the hippocampus occur during quiet immobility and slow-wave sleep (Bragin et al., 1999b; Bragin et al., 1999a; Csicsvari et al., 1999), and are thought to be critical for memory consolidation (Girardeau and Lopes-Dos-Santos, 2021). Extracellular voltage signals, including oscillations, are driven by currents entering (sinks) or leaving (sources) the intracellular brain compartment and reflect both synaptic activity and action potential firing (Mitzdorf, 1985; Liebe et al., 2011; Buzsaki et al., 2012; Schomburg et al., 2012; Pesaran et al., 2018). Importantly, excitatory and inhibitory neurons in the hippocampus fire at different phases of theta, gamma and ripple oscillations in a precise, subtype-specific manner (Klausberger et al., 2003; Klausberger and Somogyi, 2008; Cardin, 2018), and synaptic interactions between excitatory and inhibitory cells are implicated in the generation of oscillations across the spectrum, although the details may vary between frequency bands, between different brain areas and even between cortical layers (Kopell et al., 2010; Wang, 2010; Whittington et al., 2018).

At the high end of the frequency spectrum, ultrafast, 250-600 Hz oscillations have been observed in both hippocampus and neocortex. In the hippocampus such oscillations (called “fast ripples”) are almost exclusively paroxysmal (reviewed in Gulyas and Freund, 2015; Levesque and Avoli, 2019), although their underlying cellular and network mechanisms remain under debate (Dzhala and Staley, 2004; Foffani et al., 2007; Engel et al., 2009). In the somatosensory neocortex, however, 500-600 Hz non-paroxysmal wavelets have been extensively reported in humans in response to peripheral sensory or electrical stimulation (reviewed in Curio, 2000). Similar ultrafast LFP oscillations were also reported by a handful of studies in monkeys, piglets and rats (Jones and Barth, 1999; Baker et al., 2003; Ikeda et al., 2005), but their underlying mechanisms and functional significance have remained largely unexplored.

We recorded extracellular and intracellular activity evoked in thalamorecipient layers of mouse somatosensory (barrel) cortex by ex-vivo optogenetic activation of ChR2-expressing thalamocortical axons. We report here that brief light pulses (as short as 1 ms) elicited a transient (<25 ms), highly reproducible LFP oscillation or “ripplet”, which closely resembled hippocampal ripples but, at >400 Hz, was more than twice as fast. Paired whole-cell recordings from fast-spiking (FS) inhibitory interneurons and regular spiking (RS) excitatory cells revealed precise phase relationships between FS spikes, RS spikes and ripplets, and suggested that phasic RS→FS excitation is necessary for ripplet generation. Ripplets may be a ubiquitous cortical response to exceptionally salient stimuli, and could be used by the cortex to encode specific features of the sensory stimulus. Importantly, optogenetically-elicited ripplets are a uniquely accessible model for studying the synaptic basis of fast and ultrafast neural oscillations.

## RESULTS

We recorded extracellular and intracellular responses in layer 4 (L4) of barrel cortex brain slices, evoked by widefield optogenetic activation of thalamocortical synapses. Slices were prepared from brains of 3-7 postnatal weeks old mice of either sex, expressing ChR2 in the somatosensory thalamus. We applied light pulses through the epi-illumination light path of the microscope, likely activating most of the thalamocortical axons and terminals within the local L4 barrel. To verify that ChR2 was expressed by thalamocortical and not intracortical axons, we imaged and quantified the extent of Cre-dependent tdTomato reporter expression in the cortex (Fig. 1-Figure Supplement 1). Within the barrel cortex there was a sparse population of tdTomato expressing cell bodies in L2/3 and L5, but virtually none within L4 (Fig. 1 – Source Data 1). Since L4 itself is the main source of non-thalamic excitatory input to L4 barrels (Feldmeyer et al., 1999; Petersen and Sakmann, 2000), we conclude that the excitatory responses we observed were predominantly triggered by thalamocortical axons, although we cannot rule out that a small number of corticothalamic neurons in L6, which send axonal collaterals to L4 (White and Keller, 1987; Kumar and Ohana, 2008), expressed ChR2 and provided a minor excitatory contribution.

### Optogenetic activation of thalamocortical axons evoked ripple-like extracellular waveforms

A brief (1-8, typically 2-5 ms) light pulse evoked a stereotypical field potential wavelet in L4 (Fig. 1A). In this and all other figures, the light-evoked response is represented by 25 ms-long record beginning 1 ms before light onset (except for Fig. 5 in which records are 12.5 ms long); the timing of the light pulse is indicated by a cyan bar above or below the trace. At all pulse durations, the waveform consisted of an early low-amplitude negative component (solid arrowhead), followed by 2-5 (typically 3-4) larger negative transients (hollow arrowheads) riding on a slow, positive-going envelope. In this example slice, the 4^th^ transient was barely discernible in response to a 1 ms light pulse (left), but became noticeably larger with a 5 ms pulse (right). A low-amplitude negative shoulder or “hump” (arrow) was often observed on the rising phase of the first large transient. These optogenetically evoked waveforms were reminiscent of ripples, LFP wavelets recorded from the cell body layer of area CA1 of the hippocampus; we therefore named them RIPPle-Like Extracellular Transients or “ripplets”.

**Fig. 1:**
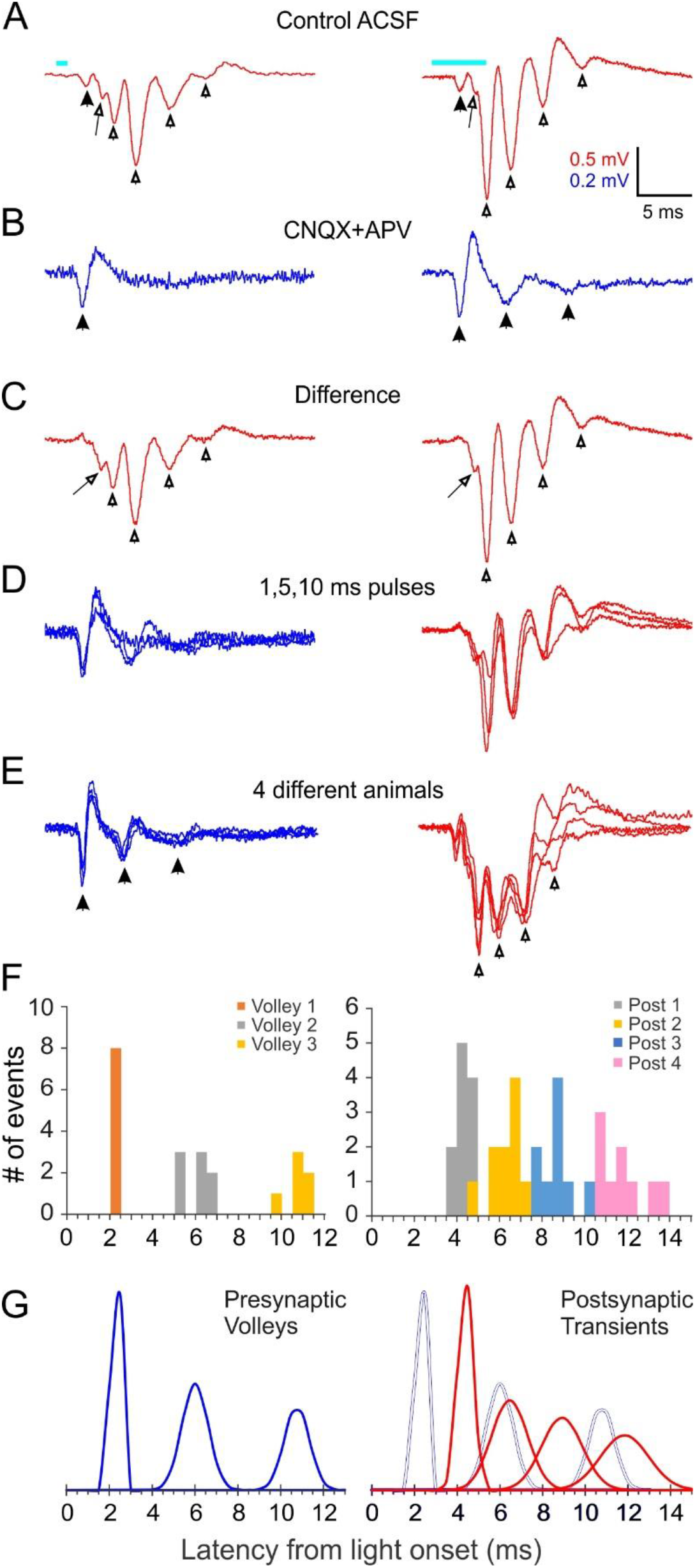
Extracellular ripplets induced by optogenetic thalamocortical activation. **A**, averaged extracellular field potential oscillations (ripplets) evoked in an example slice; light pulses (1 and 5 ms duration for left and right traces, respectively) represented by cyan bars above traces, and apply to A-C. Solid arrowhead indicates the early presynaptic TC volley; hollow arrowheads indicate the peaks of the postsynaptic components; the arrow points to the hump on the rising phase of the first postsynaptic peak. **B**, the iGluR-independent (i.e. presynaptic) component: same stimuli as in A but in the presence of CNQX+APV. Arrowheads indicate a single presynaptic volley in response to a 1 ms stimulus and 3 volleys (of decreasing amplitude and coherence) with a 5 ms stimulus. **C**, the difference between A and B, revealing the purely postsynaptic component of the ripplet. Symbols as in A. **D**, superposition of the presynaptic (blue, left) and postsynaptic (red, right) components of the ripplets at three stimulus durations. **E**, superposition of presynaptic components (blue, left) and ripplets (red, right) from slices from 4 different animals, normalized to the same maximal amplitudes to facilitate comparison of their time course. **F**, histogram of peak times (from light onset) of presynaptic volleys (left) and of postsynaptic transients (right) in slices from 8 and 11 animals, respectively (5 slices used for both plots). Presynaptic volleys peaked at 2.3±0.07, 6.0±0.22 and 10.8±0.24 ms from light onset. Postsynaptic components peaked at 4.4±0.11, 6.5±0.26, 8.9±0.30 and 11.9±0.38 ms. **G**, The histograms from F modeled as normal distributions, with the same mean, SD and integral as the corresponding component. Left, presynaptic volleys; right, postsynaptic transients (red) with presynaptic volleys overlaid as blue outlines for comparison.

**Figure 2:**
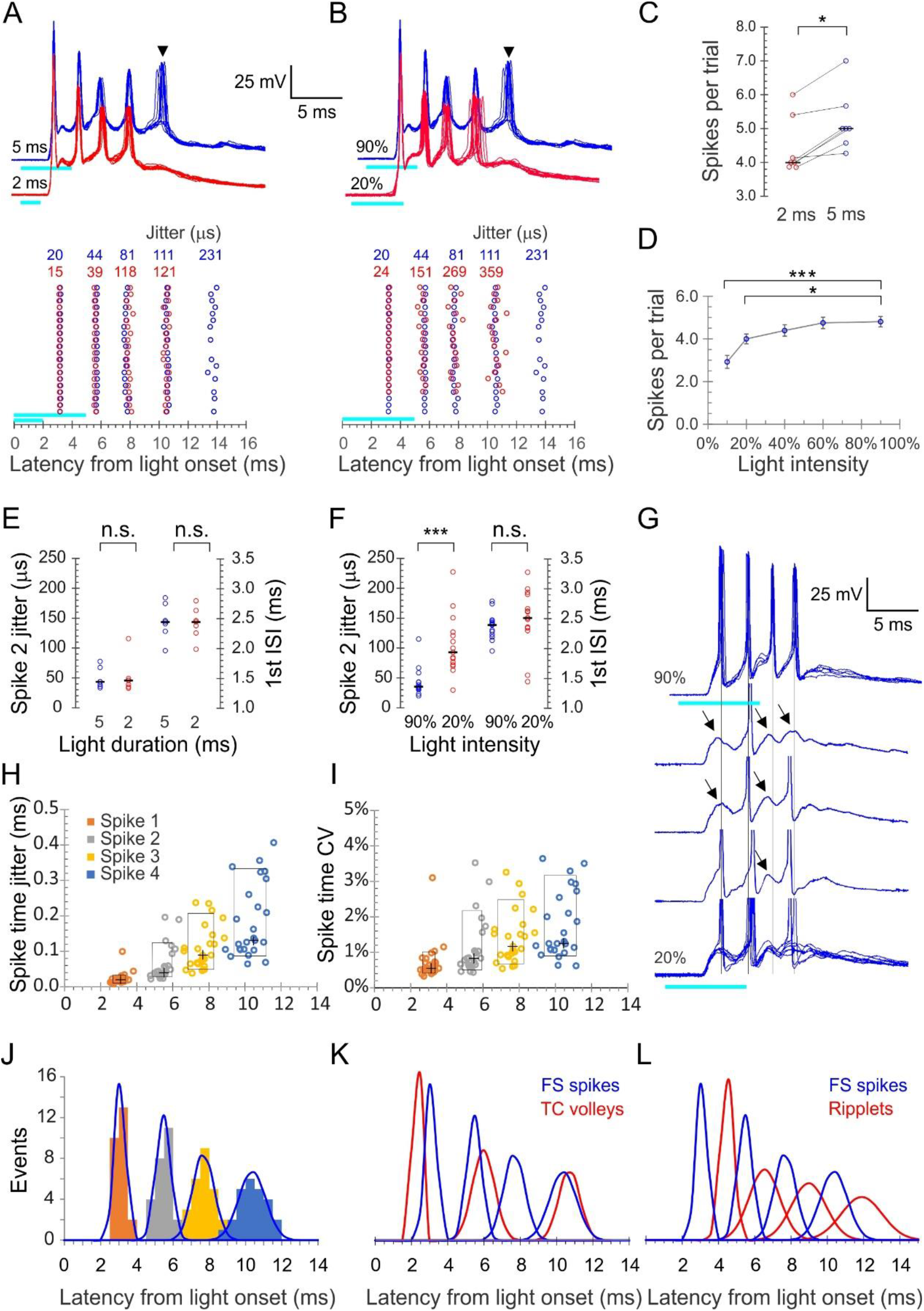
precise light-induced spike bursts in FS interneurons. **A,** upper panel, spike bursts evoked in an example L4 FS interneuron by 5 ms (blue traces) and 2 ms (red traces) light pulses (cyan bars) at 90% intensity; 10 consecutive sweeps repeated at 8 s intervals at each duration. Arrowhead indicates a 5^th^, low-reliability spike evoked by the longer light pulse. Lower panel, raster plots of 20 consecutive bursts from the same neuron, evoked by 2 and 5 ms pulses (red and blue symbols, respectively). Spike time jitters (SD of spike peak times), in μs, are indicated above the raster, in the respective colors. **B**, as in A but for bursts evoked by 20% and 90% light intensities (red and blue traces and symbols, respectively). **C**, average number of spikes/stimulus in 7 FS interneurons tested with both 2 and 5 ms light pulses. Medians indicated by horizontal black lines. *, p=0.016, sign test. **D**, the number of spikes/stimulus fired by 14 FS interneurons tested at 4-5 different light intensities. ***, p<0.0001; *, p=0.02. **E**, jitter of the 2^nd^ spike in the burst, and the 1^st^ ISI, compared between 5 and 2 ms light pulses, from the dataset of panel C; n.s.=not significant. **F**, same as E, but comparing the lowest and highest light intensities from the dataset of panel D. Note significant difference in spike jitter (***, p<0.0001) but not ISI. **G**, another example cell stimulated at 90% intensity (upper trace, 5 superimposed sweeps) and at 20% intensity (lower traces, 3 single traces and 8 superimposed sweeps). Note that at 20% intensity, spikes drop out sporadically and reveal subthreshold EPSP (arrows), with little change in the temporal structure of the burst (vertical guidelines are aligned with the spikes at 90% intensity). **H**, Spike times in all 25 FS cells with each spike order in a different color, plotted by its mean time measured from light onset (X axis) and by its jitter (Y axis); crosses indicate medians, boxes indicate 10-90^th^ percentile range. **I**, as in H but plotted along the Y axis by CV (jitter/mean spike time). **J**, histogram of all average spike times using the same color scheme as in H; each peak in the histogram is overlaid by a gaussian with the same mean, SD and integral (blue curves). **K**, The gaussians from panel 2J, representing FS spike bursts, and from fig. 1G, representing thalamocortical spike volleys, are superimposed (blue and red curves, respectively). **L**, gaussians representing FS spike bursts (blue) and ripplets (red) superimposed; note the anti-phase relationship between FS spikes and ripplets.

**Fig. 3:**
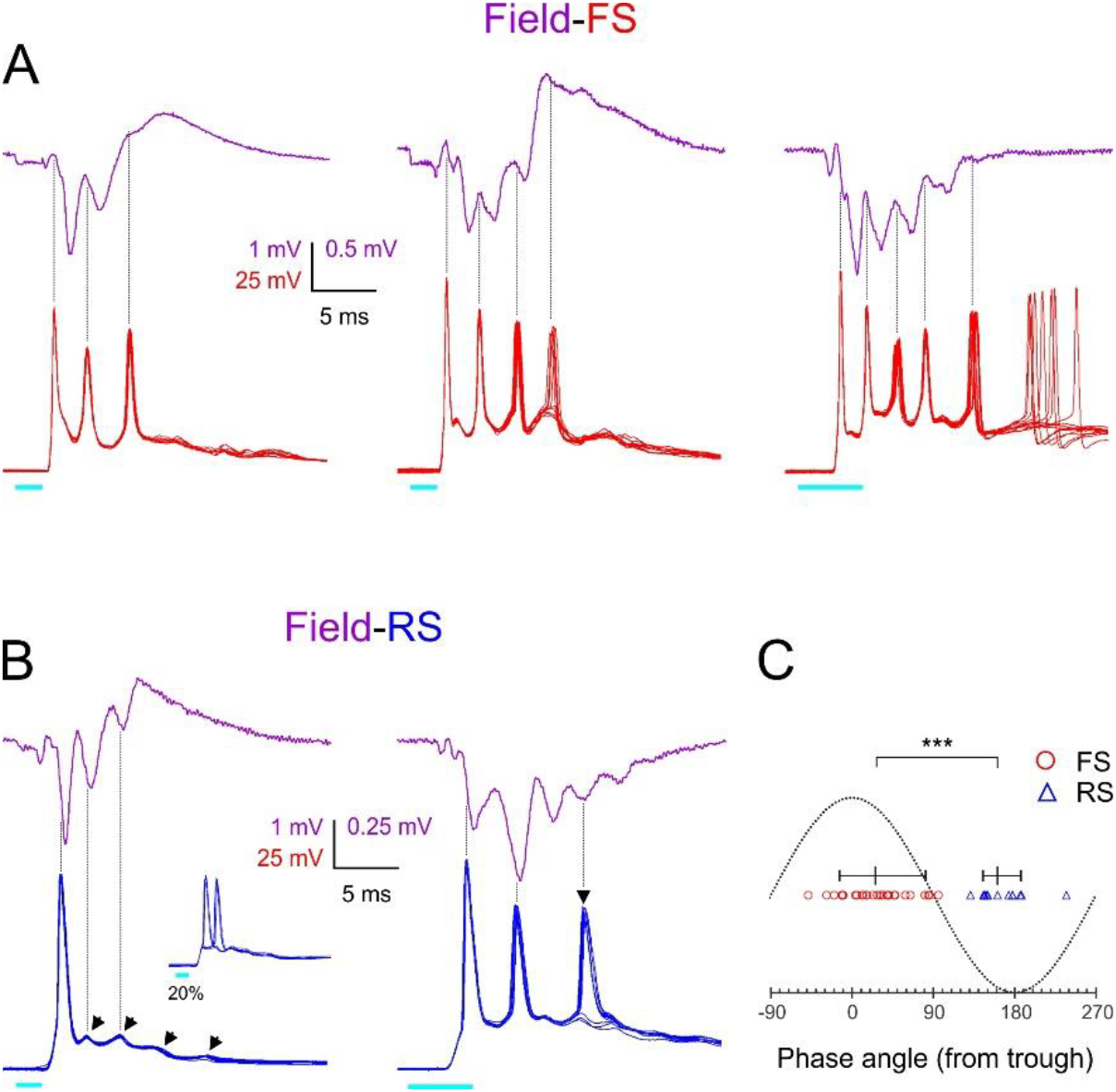
Phase-locking of FS and RS spikes to ripplets. **A**, averaged ripplets (upper traces, purple) and simultaneously recorded FS spike bursts (lower traces, superimposed consecutive sweeps, red) in response to 2 or 5 ms light pulses (cyan bars) in slices from 3 different animals. Vertical guidelines are aligned with FS spike peaks; note that FS spikes aligned with ripplet troughs. The leftmost two slices were not tested in TTX, and therefore the square-wave stimulus artifact is not subtracted and partially occludes the presynaptic volley. **B**, as A but lower traces are from 2 RS cells. Arrowheads point to subthreshold EPSPs. Note that RS spikes and EPSPs were aligned with ripplet peaks. Inset in left panel illustrates the same cell stimulated at a weaker light intensity, with the spike “hopping” between the 1^st^ and 2^nd^ EPSPs. **C**, a phase plot with ripplet troughs and peaks designated as 0^0^ and ±180^0^, respectively. FS and RS spike peaks are indicated by red circles and blue triangles, respectively. Horizontal lines with tick marks indicate medians and 10-90^th^ percentile ranges, respectively. ***=p<0.0001.

**Fig. 4:**
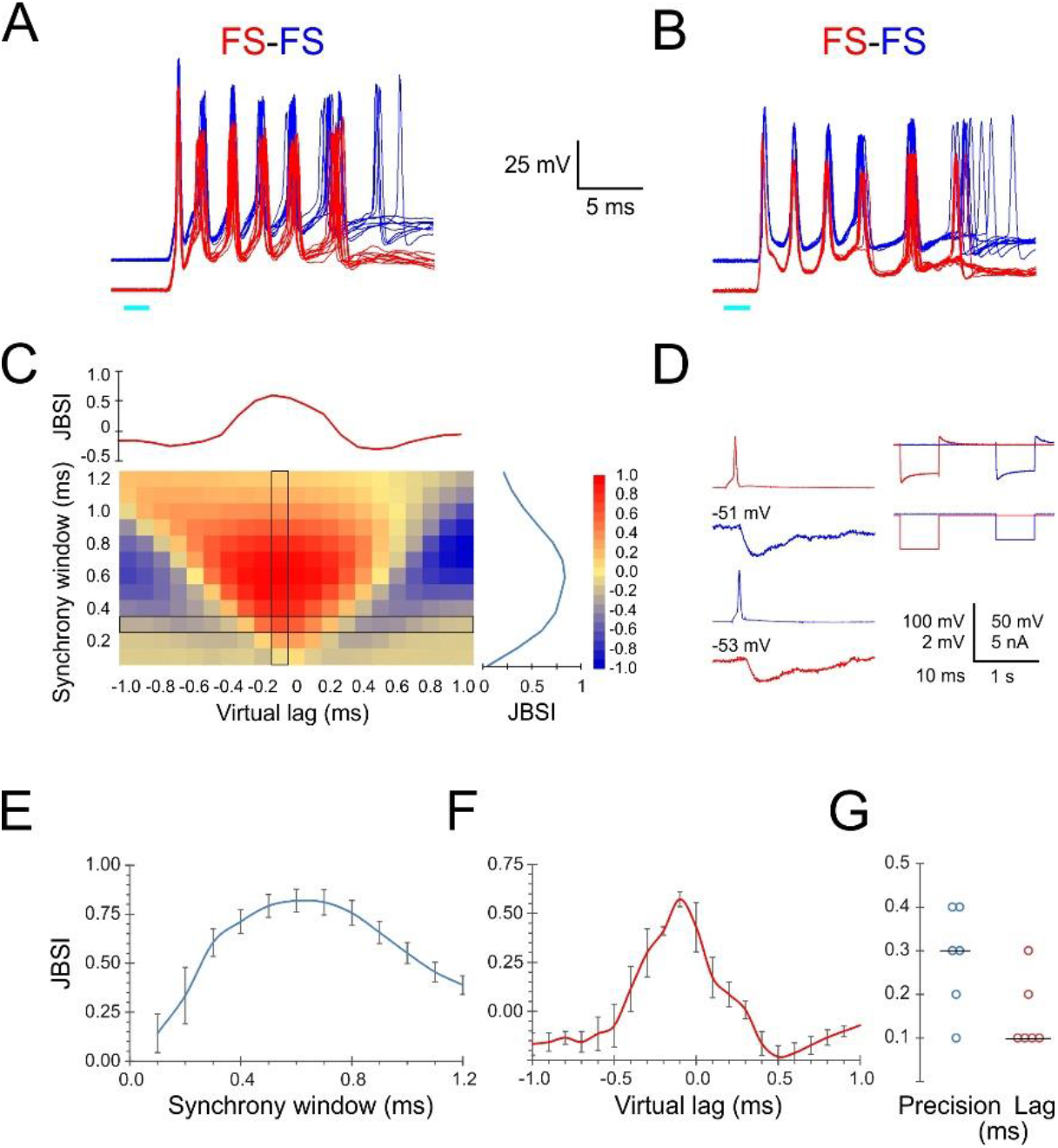
Precise firing synchrony between L4 FS cells. **A, B**, simultaneously recorded spike bursts (8 superimposed sweeps each) from two example FS-FS pairs. Traces from the two cells are displaced vertically for clarity. **C**, heat map-coded matrix of jitter-based synchrony index (JBSI) values calculated from ~300 spikes from each FS cell in A, for a range of synchrony windows (vertical axis) and virtual lags (horizontal axis) in 0.1 ms increments. The plots to the right and above the matrix are cross-sections along the column and row, respectively, indicated by a black border. The matrix element at the intersection corresponds to the smallest synchrony window with JBSI > 0.5 and the highest JBSI value along that row, and was used to define the pairwise precision and lag of this pair. **D**, averaged traces of reciprocal unitary IPSPs between the two FS cells in B (left), with holding potentials indicated, and a test for electrical coupling (right); coupling coefficient was 0.8%. **E**, average ±SEM of vertical cross-sections (as illustrated in C) through the JBSI matrices of 6 FS-FS pairs. **F**, same for horizontal cross sections. **G**, summary of pairwise precision and lag for the 6 FS-FS pairs; horizontal lines indicate medians.

**Fig. 5:**
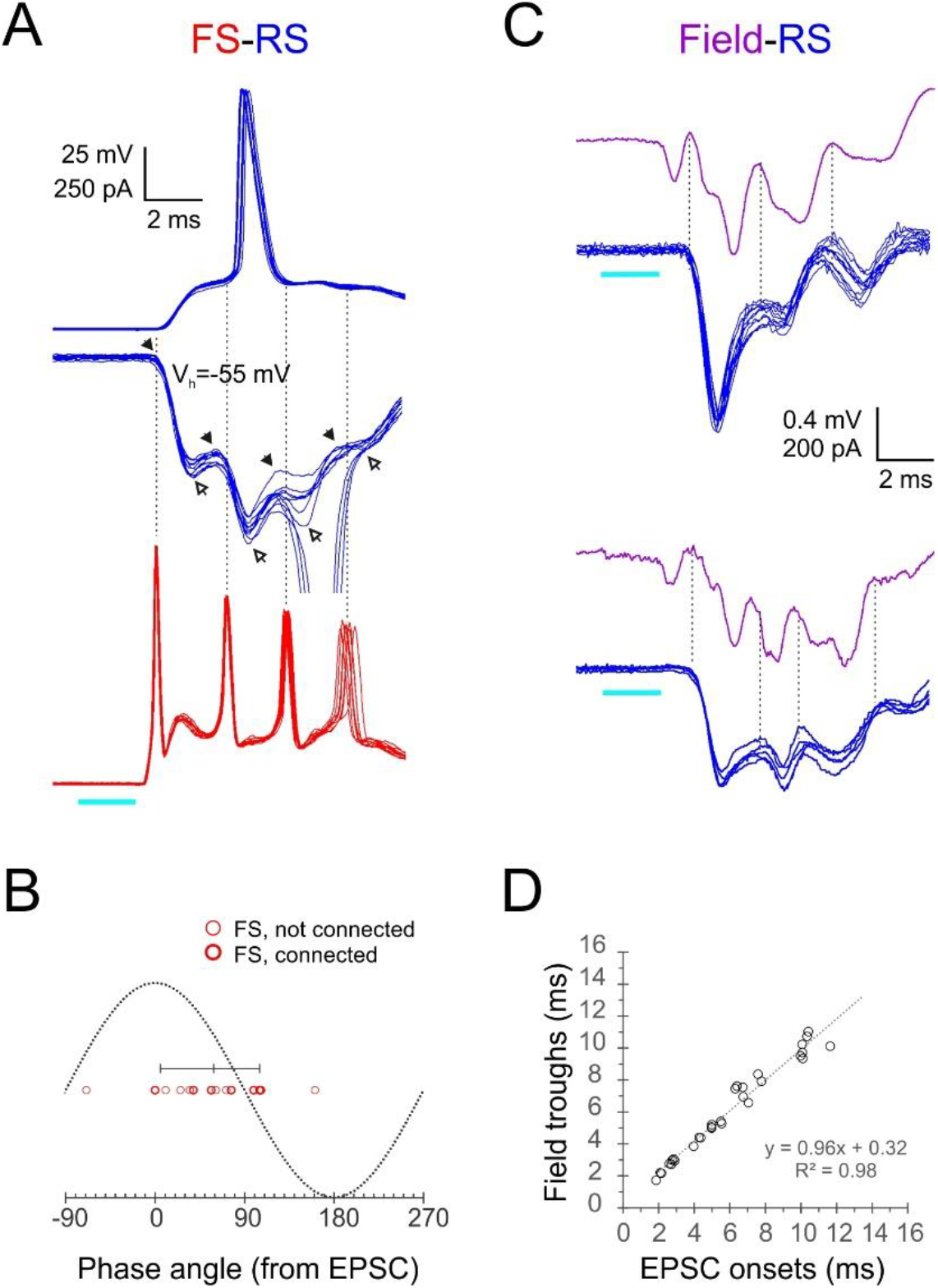
phase-locking of E/I sequences in RS cells to FS spikes and ripplets. **A**, Superimposed FS spike bursts (lower traces, red), simultaneously recorded voltage-clamped RS EPSC-IPSC sequences (middle traces, blue), and sequentially recorded current-clamp responses from the same cell (upper traces), evoked by a 2 ms light pulse. Solid and empty arrowheads in voltage-clamp traces point to EPSC and IPSC onsets, respectively. Vertical guidelines are aligned with FS spike peaks and illustrate that FS spikes always followed an EPSC and preceded an IPSC in the RS cell. Note occasional unclamped RS spikes. **B**, a phase plot of FS spikes relative to the EPSC-IPSC oscillation in RS cells, with EPSC and IPSC onsets designated as 0° and 180°, respectively. Phase angles of FS spikes (16 spikes from 6 pairs) are indicated by red circles, with heavier symbols corresponding to pairs with a direct FS→RS inhibitory connection; vertical bar indicates median and 10-90^th^ percentile range. **C**, two example voltage-clamped RS traces (blue) recorded simultaneously with the field potential in the same barrel (purple, averaged). Vertical guidelines illustrate the correspondence between EPSC onsets in the RS cell and troughs in the oscillatory field potential. **D**, a plot of EPSC onset times vs field potential trough times; equation and R^2^ value of regression line indicated.

Synchronous spike volleys in thalamocortical terminals generate relatively large current sinks in layer 4, and these are discernible as negative transients in the LFP (Morin and Steriade, 1981; Agmon and Connors, 1991; Swadlow et al., 2002; Bruno et al., 2003). To reveal any presynaptic thalamocortical components in the ripplets, we blocked fast ionotropic glutamate receptors (iGluRs) pharmacologically with CNQX+APV. This eliminated the larger negative transients but left unchanged the early negative component (Fig. 1B, left panel, solid arrowhead) which therefore reflected the initial thalamocortical volley. Interestingly, the presynaptic response to 5 ms or longer light pulses revealed one or two additional presynaptic transients of decreasing amplitude (Fig. 1B, right panel, solid arrowheads), indicating that some thalamocortical axons fired additional volleys of decreasing synchrony. Subtracting the waveforms in CNOX+APV from those in control ACSF isolated the postsynaptic component of the response (Fig. 1C). Superposition of the isolated presynaptic (Fig. 1D, left) and postsynaptic (Fig. 1D, right) contributions at 1, 5 and 10 ms stimulus durations demonstrated that the longer light pulse added late components to the response waveform but did not affect the timing of the earlier peaks. To confirm that the iGluR-independent signals reflected presynaptic action potentials we added TTX to the bath, which blocked all remaining responses except for a small square waveform. The latter was coincident with the light pulse and was most likely an electrical or optoelectrical artefact, and is subtracted from all traces in Fig. 1.

Ripplet waveforms, as well as their isolated presynaptic components, were remarkably consistent between animals in their temporal structure, as illustrated in Fig. 1E by the superposition of response components in slices from 4 different animals, drawn at different vertical scales to facilitate time course comparison. To quantify the variability in waveforms between animals, we measured the peak times (relative to light onset) of the presynaptic volleys (Fig. 1F, left; 8 slices from 8 animals tested in CNQX+APV) and of the postsynaptic components (Fig. 1F, right; 11 slices from 11 animals with ripplet amplitudes ≥0.5 mV). Standard deviations (SDs) of the presynaptic volleys ranged from 0.2-0.6 ms, and SDs of the postsynaptic components from 0.3-1.0 ms. Calculated over the 3 volleys, the presynaptic axons appeared to fire at an average frequency (=1/average ISI) of 239±6 Hz. Calculated over the 4 postsynaptic peaks, ripplet frequency averaged 408±15 Hz, i.e. nearly twice as fast as the presynaptic axons. To facilitate time course comparison of pre- and postsynaptic responses we modeled the histogram peaks as Gaussians with the same mean, SD and integral as that of the respective event (Fig. 1G). Overlaying the presynaptic and postsynaptic Gaussians (Fig. 1G, right) highlighted the temporal incongruence between the two sets of events, indicating that ripplets were not a direct, 1:1 response to bursts in thalamocortical axons.

### Optogenetic stimulation of thalamocortical axons evoked precisely timed FS spike bursts

Since ripplets were not driven by presynaptic bursts, they most likely reflected network activity within L4; but in which cells? To answer this question, we examined light-induced whole-cell responses in inhibitory fast-spiking (FS) interneurons and excitatory, regular spiking (RS) cells, two subclasses known to receive direct thalamocortical inputs (Agmon and Connors, 1992; Gibson et al., 1999; Porter et al., 2001; Bruno and Simons, 2002; Gabernet et al., 2005; Sun et al., 2006; Cruikshank et al., 2007; Shigematsu et al., 2019). These two subtypes are readily identifiable in slice recordings by their firing patterns and also fall into distinct, non-overlapping clusters in post-hoc analysis of their electrophysiological parameters (Fig. 2 - Figure Supplement 1; Fig. 2 – Source Data 1).

A brief (1-10 ms, typically 2-5 ms) light pulse at 60-90% maximal intensity elicited a strong (10-30 mV) depolarization in L4 FS cells which, in 80% of cells (N=44 cells from 35 animals), culminated in a stereotypical burst of 2-7 (typically 3-5) spikes. Representative bursts in an example FS cell, evoked by 2 and 5 ms light pulses, are illustrated in Fig. 2A,B. Spiking never persisted beyond the illustrated 25 ms time window, although the underlying subthreshold depolarization often lasted for 50 ms or more. We performed detailed analysis on all FS cells that consistently fired bursts of 4 or more spikes (25 cells from 20 animals). Bursts in this dataset had three distinguishing properties:

1. Very high frequency: averaged over these 25 cells, the first three interspike intervals (ISIs) were 2.4±0.05, 2.2±0.08 and 2.7±0.13 ms (mean±SEM), respectively, for an average burst frequency (=1/average ISI) of 418±9.5 Hz, not significantly different from the extracellular ripplet frequency (p=0.56).
2. Very high temporal precision: bursts in a given cell were precisely reproducible, as illustrated in Fig. 2A,B by the near-perfect registration of consecutive sweeps, the alignment of spikes in the raster plots and the very low jitter in spike times indicated above the raster plot. In the full dataset, spike jitter averaged 23±4, 63±10, 110±12 and 183±21 μs for the four spikes, which translates to coefficients of variation (CV=SD/mean) of 0.7±0.1%, 1.1±0.2%, 1.4±0.1% and 1.8±0.2%, respectively. This low jitter is also illustrated in Fig. 2H,I, in which the vertical extent of the boxed regions indicates 10-90^th^ percentiles of spike time jitter and CV, respectively.
3. High animal-to-animal reproducibility: across slices from different animals, the distributions of peak spike times for each of the 4 spikes in the burst was very narrow: 3.1±0.06, 5.5±0.08, 7.7±0.12 and 10.3±0.14 ms after light onset. This is also illustrated in Fig. 2H,I, in which the horizontal extent of the boxed regions indicates 10-90^th^ percentile of spike times.

To examine if burst patterns were dependent on stimulus parameters (light duration and intensity), we first compared bursts elicited by 2 and 5 ms light pulses. In the example cell in Fig. 2A, a 2 ms stimulus elicited a precisely reproducible 4-spike burst (10 superimposed red traces); increasing stimulus duration to 5 ms left the timing and jitter on these 4 spikes nearly unchanged but occasionally elicited a 5^th^, long-latency spike (arrowhead) with considerably higher jitter compared to the first 4 spikes (blue traces; note increased jitter in the raster plot in the lower panel). We refer to such spikes, with higher jitter and occasional failures, as “low reliability spikes”. All 7 FS cells tested at both stimulus durations fired more spikes in response to a 5 ms compared with 2 ms stimulus, with a median increase of 1 spikes/stimulus (Fig. 2C) but with no appreciable change in timing and precision of the earlier spikes, as seen by the identical median values of the 2^nd^ spike jitter and the 1^st^ ISI at the two stimulus durations (Fig. 2E). We then compared the number and precision of spikes in response to different light intensities. Bursts elicited by 20% and 90% intensity, in the same example cell from fig. 2A, are illustrated in fig. 2B. At the lower intensity there was again one less spike, but in addition the jitter of all spikes except the first was >3× higher. In a subsample of 14 cells tested with 5 ms light pulses at 5 different intensities (10, 20%, 40%, 60% and 90% of maximal intensity), the average number of spikes/trial increased from 2.9 at 10% to 4.8 at 90% intensity (Fig. 2D). As seen by comparing the 2^nd^ spike jitter and 1^st^ ISI between at 20% and 90% light pulses (Fig. 2F), overall spike timing did not differ between the two intensities (p=0.38) but at lower intensities spikes were much less precise (2nd spike jitter:106±13 vs. 43±6 μs, p<0.0001). That ISIs were largely independent of stimulation intensity suggested that the burst pattern was not intrinsically generated in the FS cell but was imposed on it by extrinsic inputs. Consistent with this conclusion, reducing the light level resulted in some cells in spikes dropping out sporadically, revealing apparent subthreshold EPSPs which remained aligned with the temporal structure of the burst (Fig. 2G, guidelines). That EPSPs elicit FS spikes with such high precision is consistent with the properties of excitatory synaptic inputs onto FS interneurons in neocortex and hippocampus (Geiger et al., 1997; Galarreta and Hestrin, 2001).

What were the temporal relationships between FS spike bursts, thalamocortical volleys and ripplets? We plotted all FS spike times in our dataset as a histogram and then modeled each spike order as a Gaussian (Fig. 2J). Overlaying Gaussians representing FS spikes with those representing thalamocortical volleys from Fig. 1G (Fig. 2K) illustrated the close temporal relationship between the first FS spike and the initial thalamocortical volley, with the postsynaptic spike following the presynaptic volley at a latency of 0.8 ms, consistent with a monosynaptic response. Later FS spikes, however, actually preceded the closest presynaptic volley, consistent with the conclusion above that the presynaptic bursts did not drive the postsynaptic bursts in a 1:1 manner. In contrast, overlaying the distributions of FS spikes and ripplet peaks (Fig. 2L) revealed a 1:1 relationship between these events, however with the two oscillations almost exactly out-of-phase: each ripplet peak lagged behind the corresponding spike peak by 0.44-0.54 of a cycle, translating to 1.0-1.3 ms.

### FS and RS spikes were phase-locked to the ripplet oscillation

To examine directly and in more detail the temporal relationship between FS spike bursts and ripplets, we recorded light-evoked bursts in 9 FS cells (from 9 animals), simultaneously with the LFP evoked in the same barrel. Three example from different animals are illustrated in Fig. 3A, with averaged ripplets (purple traces) and 10 superimposed intracellular FS bursts (red traces). As indicated by the vertical guidelines, FS spike peaks were closely aligned with the troughs in the extracellular waveforms, almost exactly out-of-phase with ripplet peaks, consistent with the conclusion above from the comparison of the population distributions in Fig. 2L. Fig. 3A also illustrates that the number of FS spikes in the different slices (3, 4 and 5, respectively) was exactly one more than the number of ripplet peaks (2, 3 and 4), reflecting the fact that a spike occurred in every trough, from the one preceding the first to the one following the last ripplet peak.

We also recorded light-evoked responses in excitatory RS neurons, which are the majority cell type in L4. RS cells responded to the light stimulus with a large depolarization (up to 15-20 mV above resting potential), crested by a series of 3-5 apparent EPSPs, typically giving rise to 1-2 spikes (Fig. 3B, blue traces). Of 55 RS cells recorded in current clamp mode and stimulated at a near-maximal light level, 73% fired at least one, but never more than 2 spikes/stimulus; the rest either remained subthreshold (11%) or occasionally fired 3 or more spikes (16%). Spikes evoked by near-maximal stimulus levels were mostly highly precise (Fig. 3B left, 90% intensity), but spikes arising from late EPSPs, or spikes evoked by lower light levels, were often low-reliability spikes which occasionally failed (Fig. 3B right, arrowhead) or which sporadically switched between EPSPs, a pattern we refer to as “EPSP hopping” (Fig. 3B left, inset). To examine phase relationships between RS spikes and ripplets, we recorded in current-clamp mode from 10 RS cells (from 10 animals) simultaneously with the LFP. As indicated by the vertical guidelines in the two example records in Fig. 3B, peaks of RS spikes and EPSPs occurred near the field potential negative peaks, i.e. in-phase with ripplets. This was reminiscent of hippocampus CA1 ripples, in which pyramidal cells fire close to ripple peaks (Ylinen et al., 1995; Klausberger et al., 2003).

We quantified the phase relationships between ripplets and FS and RS spikes in these two datasets (of 9 and 10 cells, respectively) by assigning each ripplet trough a phase angle of 0°, and assigning phase angles of −180° and 180° to the negative peaks preceding and following the trough, respectively. We then assigned a phase angle to each of the 28 FS spikes and 13 RS spikes observed in these experiments, by interpolating its time of peak between the nearest ripplet trough and peak (Fig. 3C). The phase angles of the two cell types were non-overlapping, with FS spikes occurring at a median phase angle of 26° (i.e. slightly past the trough; 10-90^th^ percentile range: (−14) −82°, translating to a range of 0.65 ms), and RS spikes occurring at a median phase angle of 161° (i.e. slightly before the peak; 10-90^th^ percentile range: 145 – 187°, translating to a range of ~0.3 ms). Thus, on average, RS spikes lagged behind FS spikes by ~135° or ~ 0.9 ms. Given the 2.4 ms ripplet cycle period, this also means that RS spikes occurred ~1.5 ms before the following FS spike. A 1.5 ms lag is just enough for RS cells to generate monosynaptic EPSPs in FS cells, and for postsynaptic FS cells to reach threshold and fire (Galarreta and Hestrin, 2001).

### FS cells synchronized their firing with near-zero lag and submillisecond precision

If FS spikes are phase-locked to the field potential in their local barrel, one would expect different FS cells in the same barrel to fire in tight synchrony. We therefore examined intra-barrel FS-FS temporal relationships by simultaneous paired recordings. Indeed, when stimulated by a light pulse, simultaneously recorded FS-FS pairs exhibited highly precise spike synchrony. In the two example pairs shown in Fig. 4A, B, the peaks of the first three spikes in each burst aligned to within 0.1-0.3 ms between the two cells. To quantify the magnitude and precision of pairwise synchrony, we used the Jitter-Based Synchrony Index (JBSI), which is a measure of spike-spike synchrony in excess of that expected by chance (Agmon, 2012) (see Methods). Synchrony is defined as the fraction of spikes co-occurring within a pre-determined “synchrony window” (SW), and chance synchrony is the average synchrony remaining after shifting each spike in one train by a random jitter ≤J, J=2·SW. When calculated for decreasing SW values, the JBSI typically increases to a maximum and then precipitously drops off to zero, as J falls below the intrinsic precision of the system and the applied jitter no longer disrupts the pairwise synchrony. We defined “pairwise precision” as the smallest SW value with JBSI>0.5. Since neurons can maintain a precise temporal relationship even when they fire at non-zero time lags relative to each other, one can extend the analysis by calculating the JBSI for a range of virtual lags (shifts) of one spike train relative to the other, analogous to a classical cross-correlation histogram; we defined “pairwise lag” as the virtual lag for which the JBSI was maximized. The full analysis can be represented as a 2-dimensional matrix in which the JBSI is plotted as a function of the two parameters, SW and virtual lag, as illustrated in Fig. 4C. Vertical and horizontal cross-sections through the matrix are plotted to the right and top of the matrix, illustrating how the JBSI varied with SW and virtual lag values, respectively.

We calculated the JBSI matrices for 6 FS-FS pairs from which at least 100 spikes were recorded per cell. The averaged horizontal and vertical cross-sections through the 6 matrices are plotted in Figs. 4E,F, respectively, and the pairwise precision and lag (in absolute value) of the 6 pairs are plotted in Fig. 4G. The median pairwise precision and lag were 0.3 and 0.1 ms, respectively. While there is some degree of arbitrariness in the exact definition of pairwise precision, this analysis indicates that FS spikes within a given barrel synchronized with near-zero lag and with sub-millisecond precision.

What accounted for this highly precise FS-FS synchrony? Precise spike synchrony is most often attributed to electrical coupling via gap junctions (Galarreta and Hestrin, 1999; Mancilla et al., 2007), to inhibitory synaptic connectivity (Gibson et al., 2005; Hu et al., 2011) or to common excitatory inputs (Perkel et al., 1967; Sears and Stagg, 1976; Alonso et al., 1996). However, unlike rat cortex, coupling between FS interneurons in the mouse is weak (average coupling coefficient of ~1.5% (Galarreta and Hestrin, 2002; Meyer et al., 2002; Hu et al., 2011; Hu and Agmon, 2015). The right panel in Fig. 4D illustrates how we tested for electrical coupling, which in all 6 pairs was absent or barely detectable (coupling coefficient <1%). In addition, inhibitory synaptic connections were observed in only 2 of the 6 FS-FS pair, including the pair of Fig. 4B (Fig. 4D, left panels) but not the pair of Fig. 4A. We conclude by elimination that the precise FS-FS synchrony we observed was likely driven by common excitatory inputs.

### FS spikes and ripplet troughs were phase-locked to EPSC-IPSC sequences in RS cells

As both FS and RS cells were phase-locked to the ripplet oscillation, albeit at different phases (Fig. 3C), they would be expected to phase-lock with each other. To directly examine the temporal relationships between FS and RS cell responses to optogenetic thalamocortical activation, we performed simultaneous recordings from 6 FS-RS pairs (from 4 animals) with the RS cell alternating between current and voltage clamp to record both spikes and postsynaptic currents. When in voltage clamp, we held the RS cells at −50 to −55 mV to reveal both EPSCs (negative deflections) and IPSCs (positive deflections). We also tested recorded pairs for direct synaptic connections between them.

An example pair is shown in Fig. 5A, with 10 superimposed FS spike bursts (red) in the lower traces, simultaneously recorded RS PSCs in the middle traces (blue) and sequentially recorded RS spikes (blue) in the upper traces. The voltage-clamped RS responses revealed a sequence of regularly alternating EPSCs and IPSCs (E’s and I’s, marked by filled and hollow arrowheads, respectively). The dashed vertical guidelines are aligned with FS spikes peaks, and demonstrate that each FS spike occurred immediately following the onset of an EPSC and before the onset of an IPSC in the RS cell. IPSCs followed FS spikes whether or not the FS interneuron directly inhibited the RS cell. To quantify this temporal relationships we assigned a phase angle to each of the 16 FS spikes observed in this dataset, by defining RS EPSC onsets as 0° and RS IPSC onsets as 180° (Fig. 5B). (Note that in these experiments light intensity was reduced to minimize unclamped RS spikes, thereby also reducing the number of FS spikes). The median FS spike phase angle was 59° (10-90^th^ percentile range: 5-106°). Given a 2.4 ms oscillation period, each FS spike occurred, on average, 0.4 ms after the onset of an EPSC in the RS cell and 0.8 ms before the onset of an IPSC in the same RS cell. The most parsimonious interpretation for these temporal relationships is that RS and FS cells received a barrage of common EPSPs at the ripplet frequency, these EPSPs elicited short-latency (0.4 ms) spikes in the FS cells and these spike volleys, in turn, elicited IPSPs in RS cells after a brief (0.8 ms) synaptic delay. Given that RS spikes occurred, on average, 0.9 ms after the previous FS spike volley (Fig. 3C), this delay indicates that spikes in excitatory cells occurred nearly simultaneously with the onset of IPSPs in the same cells, leaving a very narrow “window of opportunity” for RS cells to fire. The earliest cohort of RS cells to reach spike threshold fired, while spiking in other RS cells, those slightly slower to depolarize, was preempted by a strong volley of IPSPs. This explains why only a fraction of RS cells fired on any given cycle, and also explains how synchronous firing was maintained from one cycle to the next.

Given the similarity in phase distributions of FS spikes in relation to extracellular ripplets (Fig. 3C) and to E-I sequences in RS cells (Fig. 5B), we also examined the temporal relationships between ripplets and E-I sequences directly, by recording from 8 RS cells (in 8 slices from 4 animals) in voltage-clamp mode, simultaneously with the LFP. Two example slices are shown in Fig. 5C, with superimposed voltage-clamped records from the RS neuron (blue) below the simultaneously recorded, averaged field potential (purple); the vertical guidelines demonstrate that the troughs of the ripplet waveforms were almost exactly aligned with the onset of EPSCs in the RS cells. A plot of ripplet troughs against EPSC onsets in the full dataset demonstrates their close temporal coincidence (Fig. 5D). This close fit suggests that the negative ripplet peaks reflected current sinks generated by population EPSPs in RS cells.

### L5 FS spike bursts were driven by L4 RS cells

What was the source of the common EPSPs evoked in L4 FS and RS cells? Clearly, the initial EPSP was a response to the light-activated thalamocortical spike volley; thalamocortical axons in barrel cortex are known to make divergent connections onto both excitatory cells and fast-spiking interneurons (Gabernet et al., 2005; Cruikshank et al., 2007). It seems unlikely, however, that later EPSPs were elicited by thalamocortical volleys, as indicated above (Fig. 2K). The other main source of excitatory synapses in a L4 barrel are excitatory neurons in the same barrel, known to make synaptic contacts on each other and on L4 FS cells (Beierlein et al., 2003; Staiger et al., 2009; Koelbl et al., 2015). If EPSPs from local excitatory cells were driving FS spike bursts, then blocking or suppressing firing in L4 RS cells should eliminate or reduce the occurrence of the 2^nd^ and later spikes in each FS burst. To test this, we attempted to suppress firing in L4 excitatory cells optogenetically, by infecting them with intracranially injected AAV vectors coding for the inhibitory, red-activated chloride pump JAWS. Unfortunately, we found that expression of the CaMK2-driven construct in L4 was very weak. We observed, however, strong JAWS expression in L5 excitatory cells. Since L5B is innervated by the lower tier of thalamocortical terminals, we tested L5B FS cells with light pulses targeted to L5. L5B FS cells responded with bursts which were identical, in spike number and frequency, to those observed in L4 (3.9±0.2 spikes/stimulus at an average frequency of 417±15 Hz, N=12 cells from 10 non-injected animals). We therefore attempted to block ex-vivo bursts in L5 FS cells from JAWS-infected animals by applying 50-100 ms long inhibitory red light pulses, superimposed upon brief excitatory blue light pulses. JAWS activation completely suppressed firing in half of all L5 RS cells which fired in response to blue light alone (15 RS cells from 10 JAWS-injected animals were tested, 6 of which fired to blue light alone). Surprisingly, this suppression was accompanied by a negligible reduction in spikes/stimulus in L5 FS cells (0-6% reduction; N=4 cells from 3 animals). We hypothesized that blue light stimulation in L5 activated thalamocortical axons *en route* to L4, eliciting ripplets in L4, and that recurrent translaminar connections from L4 excitatory cells to L5 FS interneurons (Pluta et al., 2015) were driving L5 FS bursts (Fig. 6D). To test this hypothesis we recorded from a subset of slices after making a cut along the L4/L5 boundary, thus disconnecting the two layers while leaving the thalamocortical input to L5, as well as local connectivity within L5, mostly intact. As illustrated in Fig. 6B and summarized in Fig. 6C, L5 FS cells in cut slices fired, on average, 1.6±0.3 spikes/stimulus (17 cells from 9 animals), compared with 3.9±0.2 spikes/stimulus in intact slices (11 cells from the same 9 animals; p<0.0001). This effect was not attributable to decreased FS interneurons’ excitability in the cut slices, as resting potential, firing threshold and input resistance were not significantly different between groups (p>0.50 for each comparison). We conclude that thalamocortically induced bursts in L5 FS cells and, by extension, also in L4 FS cells, were driven by recurrent excitation from L4 excitatory neurons; in other words, phasic excitation by L4 excitatory cells drove ripplet-frequency FS bursts in both layers.

**Fig. 6:**
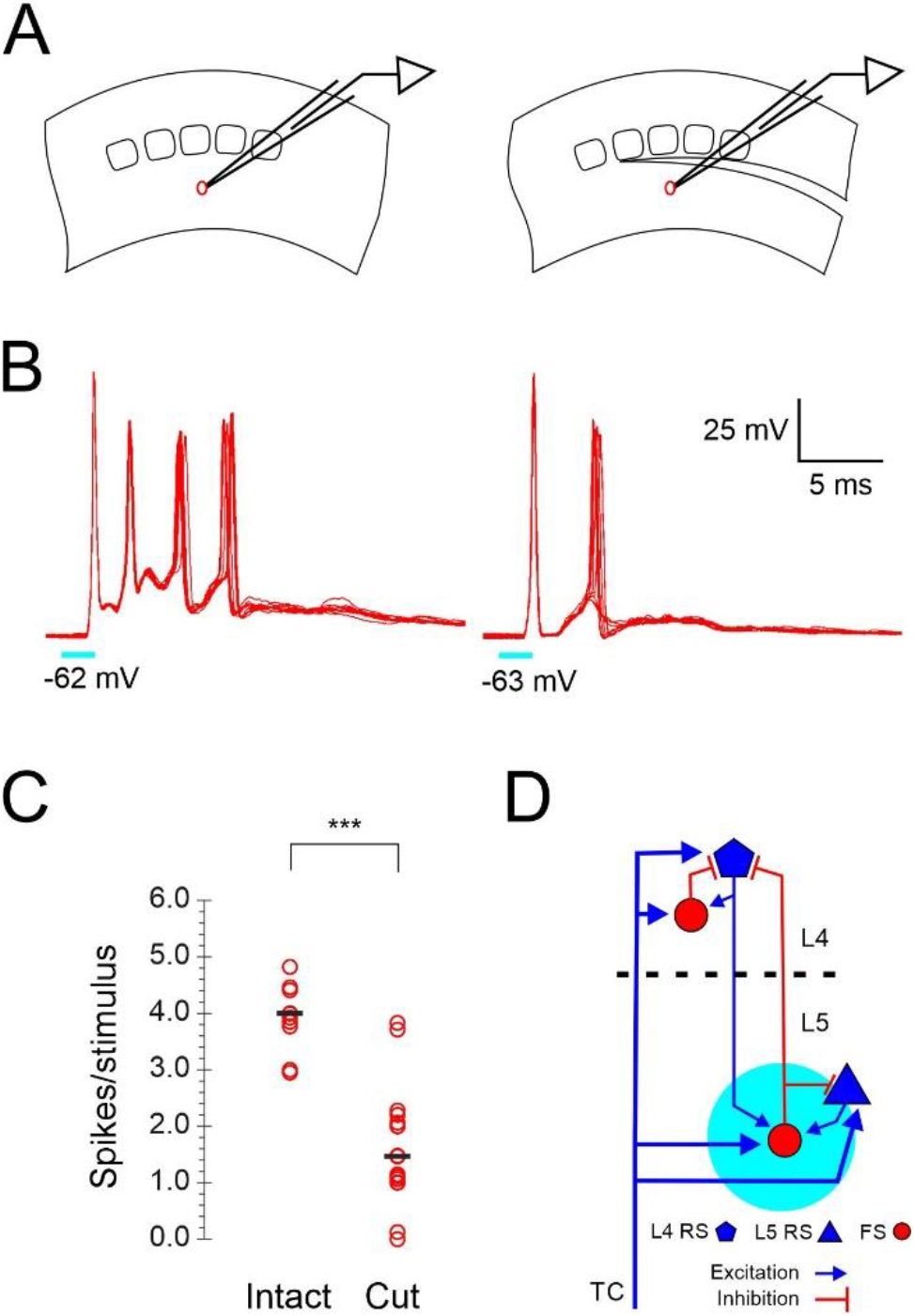
Bursts in L5 FS cells required intact L4-L5 connectivity. **A**, cartoon of the recording setup, from a L5 FS cell in an intact slice (left) and in a slice with cut through the L4/L5 border (right). **B**, superimposed spike bursts in example FS cells from an intact slice (left) and from a slice with a L4/L5 cut (right); resting potentials indicated. **C**, summary of average spike counts/ stimulus in intact and cut slices; horizontal black bars represent median values. Disconnecting L5 from L4 reduced spike counts by more than half. ***=p<0.0001. **D**, cartoon illustrating the bidirectional connectivity of L4 RS cells and L5B FS cells, which was abolished by the L4/L5 cut (dashed line). Cyan circle represents the epi-illumination light spot in L5. TC, thalamocortical axons.

## DISCUSSION

We describe here transient (<25 ms), ultrafast (>400 Hz) network oscillations, which we refer to as “ripplets”, elicited in thalamorecipient cortical layers by brief (as short as 1 ms) optogenetic stimulation of thalamocortical axons and terminals. The oscillation was reflected in the extracellular voltage (the LFP) as 2-5 negative waves (Fig. 1). In whole-cell intracellular recordings, FS cells fired a highly reproducible burst of 3-6 spikes at the same frequency as the LFP (Fig. 2), with spike peaks aligned with LFP troughs (Fig. 3A,C). As revealed by pairwise recordings, FS cells fired in exquisitely precise, sub-millisecond synchrony (Fig. 4). Excitatory RS cells rarely fired more than 1-2 spikes per ripplet, and these spikes were temporally offset from FS spikes (following them by ~0.9 ms) and aligned with ripplet peaks (Fig. 3B,C). When recorded in voltage-clamp mode at an intermediate holding potential, RS neurons were seen to receive an alternating sequence of EPSCs and IPSCs, closely in-synch both with FS spike bursts (Fig. 5A,B) and with the extracellular oscillation (Fig. 5C,D). Each FS spike followed an RS EPSC at short (0.4 ms) latency, and preceded an RS IPSC by 0.8 ms, suggesting that FS spikes were elicited by EPSPs shared with the excitatory cells, and in turn evoked monosynaptic IPSPs in the same excitatory cells.

Taken together, these observations suggest the following synaptic mechanisms for ripplet generation. The initial synchronous thalamocortical volley sets in motion a cascade of volleys or “spike packets” in L4 excitatory RS cells, each packet triggering the next one (Fig. 7A), similar to feedforward “synfire chains” (Abeles, 1991; Diesmann et al., 1999; Mehring et al., 2003; Barral et al., 2019). The 2.4 ms period of the oscillation is determined by the sum of the synaptic delay, from spike to EPSP onset, and the rise-time, from EPSP onset to spike, which in RS cells are both relatively slow. The initial thalamocortical volley, as well as each subsequent RS spike packet, also elicit highly synchronous FS spike volleys but at a shorter delay of ~1.5 ms, because synaptic delays, EPSP rise times and spikes are all faster in FS cells (Hu et al., 2014; Tremblay et al., 2016). This leaves a gap of 0.9 ms between the FS spike volley and the next RS volley, which is just enough time for FS cells to elicit monosynaptic IPSPs in RS cells (Fig. 7B). The arriving IPSP wavefront coincides with the rising phase of the RS spikes, too late to block the earliest spikes but in time to preempt any “laggard” RS cells from firing. In this manner, FS cells enforce RS firing synchrony and thereby maintain the synchrony of the next FS spike volleys. On any given cycle of the oscillation, only a fraction of all excitatory cells escape this feedforward inhibitory control and fire, enough to elicit the next cycle but not enough for the excitatory feedforward cascade to turn into a paroxysmal chain reaction. The number of excitatory cells firing declines with each cycle, likely because of build-up of inhibition, until the oscillation dies out.

**Fig. 7:**
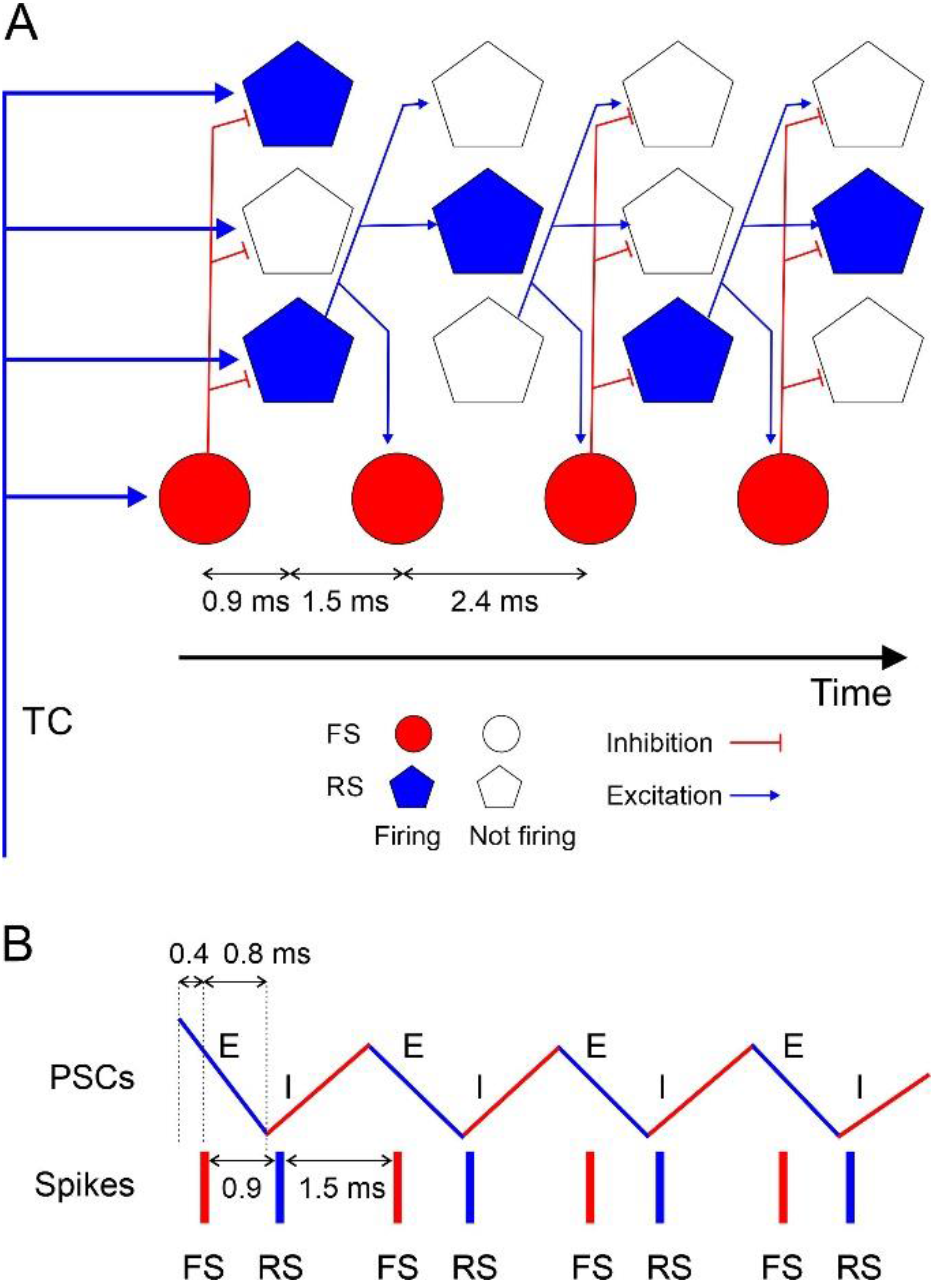
Model explaining how a ripplet oscillation arises. **A,** circuit diagram illustrating one FS interneuron and 3 excitatory RS cells at 4 successive time points, from left to right, representing the 4 cycles of the oscillation. Filled symbols represent cells firing, and their position along the horizontal axis represent spike time; hollow symbols represent cells which remain subthreshold. For clarity, only connections from one RS cell are depicted. A synchronous thalamocortical (TC) volley excites FS and RS cells in parallel; FS cells fire ~0.9 ms earlier because of the faster kinetics of their EPSCs and spikes. RS cells generate an excitatory feedforward cascade of successive spike volleys which elicit additional FS spike volleys. While FS cells fire on every cycle, RS cells typically fire on only 1-2 cycles per ripplet. The cycle period (2.4 ms) is the sum of the synaptic delay and the postsynaptic time-to-spike in RS cells. **B**, a schematic timeline of the major events during a ripplet, including FS spikes (red bars), RS spikes (blue bars) and EPSC-IPSC sequences in RS cells (zig-zag line, E’s and I’s represented by blue and red segments, respectively.) FS cells receive EPSPs shared with RS cells; they fire 0.4 ms after EPSP onset and, in turn, elicit IPSPs in RS cells after a 0.8 ms delay, resulting in coincidence of IPSP arrival with onset of spikes in RS cells, enforcing firing synchrony on RS cells.

According to this model, both phasic inhibition and phasic excitation, reverberating at the same ultrafast frequency but out of phase with each other (Fig. 7B), are necessary for generating this network oscillation. While FS cells are a critical component of the circuit, they do not act as independent pacemakers, and direct FS-FS interactions, chemical or electrical, may not be a necessary feature (see below). In contrast, RS→FS and FS→RS connections are critical, as suggested by our demonstration that disconnecting L4 and L5 by a cut disrupted FS bursts in L5 (Fig. 6). Several aspects of this model are at odds with currently proposed models of hippocampal ripples (Buzsaki, 2015), and this could reflect real differences between the two types of oscillations. Nevertheless, the phenomenological similarities between neocortical ripplets and hippocampal ripples (see below) suggest that optogenetically evoked ripplets may be a uniquely accessible model system for deciphering the cellular, synaptic and network mechanisms underlying other fast and ultrafast cortical oscillations, mechanisms which to-date remain elusive.

### Previous observations of ripplet-like oscillations in the neocortex

Non-invasive EEG and MEG recordings in humans have consistently identified fast, transient (~600 Hz, 10-15 ms) oscillations over primary somatosensory cortex, elicited by peripheral nerve stimulation, riding over the N20 peak which reflects the monosynaptic cortical response (reviewed in Curio, 2000; Hashimoto, 2000). A handful of animal studies have also reported similar observations. In a combined EEG and single-unit recording study in monkeys, 600 Hz oscillations were found to occur in phase with bursts or single spikes in putative pyramidal cells, in response to both electrical and tactile stimulation of the forearm (Baker et al., 2003). Electrical stimulation of the thalamus elicits 400-600 Hz LFP oscillations in cortical layers 4-6 in the rat, in phase with extracellular RS and FS spikes and bursts (Kandel and Buzsaki, 1997). Barth and coworkers report brief (~13 ms) periods of 350-400 Hz EEG oscillations over barrel cortex of both awake and ketamine-anesthetized rats, in response to transient, rapid-onset whisker displacements; similar oscillations were also observed over auditory cortex in response to sound clicks (Jones and Barth, 1999). In a follow-up intracellular study (Jones et al., 2000), the same authors report ~500 Hz FS spike bursts in response to whisker deflections, with individual spikes aligned to the troughs in the extracellular field potential. RS cells typically fired 1-2 spikes interspersed with subthreshold events which occurred at the frequency of the field potential oscillations, precisely as we observed here. Albeit sparse, these earlier in-vivo studies in are in close agreement with our ex-vivo results, supporting the conclusion that ripplets in our experiments were not an artefact of our slice preparation or of the optogenetic stimulation, but are a physiological, common and possibly ubiquitous neocortical response to a transient, strong and synchronous activation of thalamocortical inputs.

If ripplets are a ubiquitous response in sensory neocortex, why have they been reported by so few in-vivo studies? Most likely because most studies did not record LFPs or, if they did, low-pass filtered them below the frequency bandwidth necessary to resolve ripplets. Nevertheless, there are multiple examples in the literature of high-frequency bursts fired by FS interneurons in thalamorecipient cortical layers, in response to sensory or electrical stimulation. In awake rabbits, FS cells fire brief (3-7 spikes), ultrafast (>600 Hz) bursts in response to electrical stimulation of the thalamus, reminiscent of the FS bursts we describe here (Swadlow, 1995). FS cells in sedated rats fire, on average, 4 spikes/stimulus in response to high-velocity whisker deflections (Lee and Simons, 2004). In awake, head-fixed mice engaged in an object localization task, FS cells often fire short-latency, high-frequency bursts following active whisker touch (Yu et al., 2016; Yu et al., 2019). In these and similar examples, high-frequency FS bursts may have been embedded in ripplet oscillations which went unrecognized.

### Neocortical ripplets vs hippocampal ripples

There are many similarities between the ripplet events described here and ripples described in the CA1 and CA3 fields of the hippocampus (comprehensively reviewed in Buzsaki, 2015). Similar to CA1 ripples, individual excitatory cells did not fire on every ripplet cycle, although the majority of neocortical RS cells fired at least one spike per stimulus, whereas only about 10% of CA1 pyramidal cells fire during a given ripple event (Buzsaki et al., 1992; Ylinen et al., 1995). The subthreshold E-I sequences we observed in RS neurons (in both current clamp, Fig. 3B, and voltage clamp, Fig. 5A,C) resembled ripple-locked membrane potential fluctuations observed in mouse hippocampal pyramidal cells in behaving mice (English et al., 2014) or in slices (Maier et al., 2003). Also similarly to ripples (Ylinen et al., 1995; Klausberger et al., 2003), excitatory L4 cells fired at or near the negative peak of the LFP oscillation, while interneurons fired near the troughs. There is also similarity in the triggering mechanism, which in the hippocampus is a volley or burst of volleys in the main excitatory input to CA1, the Schafer collaterals, whereas ripplets were elicited by a spike volley in the main input to the cortex, thalamocortical axons. Moreover, in both systems the triggering input does not impose its own burst frequency on the target and only acts as a source of strong excitation, and the oscillation is generated de-novo in the target structure (Sullivan et al., 2011). Lastly, ripples in ex-vivo hippocampus consist of ~4 cycles/event (Maier et al., 2003; Hajos et al., 2013), identical to ripplets.

There were also some conspicuous differences between ripplets and hippocampal ripples, most notably in frequency: ripplet frequency was ~420 Hz under our ex-vivo recording conditions, about twice as fast as hippocampal ripples in-vivo (Buzsaki et al., 1992) or in slices (Maier et al., 2003; Ellender et al., 2010). What could account for this difference? One difference is the dense local excitatory connectivity in layer 4 of the barrel cortex (Feldmeyer et al., 1999; Petersen and Sakmann, 2000; Schubert et al., 2003; Kameda et al., 2012; Koelbl et al., 2015) compared with the sparsity of local excitatory connections in CA1 (Deuchars and Thomson, 1996). This difference extends to excitatory-to-inhibitory connectivity: Individual PV interneurons in the CA1 region receive only ~15% of their excitatory synapses from local CA1 collaterals (Gulyas et al., 1999; Bezaire and Soltesz, 2013), and these collaterals arborize mostly in the oriens/alveus layers where FS interneurons only have distal dendrites (Freund and Buzsaki, 1996). In contrast, PV interneurons in L4 of cat visual cortex receive 45% of their somatic and 70% of their proximal dendritic excitatory contacts from local L4 cells (Ahmed et al., 1997). In L4 of barrel cortex, connection probability between RS and FS cells (in either direction) is >40% (Beierlein et al., 2003). Thus, synchronous inputs from many excitatory neurons will summate in L4 RS and FS cells, generating a steep rise in membrane potential and a short latency to spiking, shortening the interval between cycles and thereby generating a faster oscillation.

There is no general agreement on the mechanisms generating hippocampal ripples, with several alternative models proposed in the literature, assigning different weights to pyramidal-interneuron (PYR-INT) and interneuron-interneuron (INT-INT) connections (Buzsaki, 2015). Recent experimental and computational studies favor the latter over the former, concluding that FS interneurons act as the central pacemakers of the ripple oscillation, and that tonic excitation from pyramidal neurons is needed only to keep interneurons sufficiently depolarized (Schlingloff et al., 2014; Stark et al., 2014; Gan et al., 2017; Ramirez-Villegas et al., 2018). In contrast, we propose that neocortical ripplet oscillations were driven, on a cycle-by-cycle basis, by phasic excitatory inputs, and that FS interneurons were followers, rather than leaders, in these oscillations. These two points are discussed in more detail in the following two sections.

### The role of excitatory-to-inhibitory connections

Our data point to RS→FS connections as the drivers of the ultrafast bursts in FS cells. We took advantage of the spatial separation between excitatory L4 cells and inhibitory L5B interneurons and demonstrated that bursts in L5B FS cells were abolished or truncated when layers 4 and 5 were physically disconnected. L5B FS interneurons receive strong thalamocortical input (Agmon and Connors, 1992; Tan et al., 2008; Constantinople and Bruno, 2013; Yu et al., 2019), as well as strong excitatory input from L4 (Pluta et al., 2015). Conversely, many FS interneurons straddling the L5/6 border have a substantial axonal arbor in L4 (Porter et al., 2001; Kumar and Ohana, 2008; Bortone et al., 2014; Frandolig et al., 2019; Gouwens et al., 2020). The results of the L4/5 cut experiments imply that this strong bidirectional connectivity (RS^L4^→FS^L5B^ and FS^L5B^→RS^L4^, Fig. 6D) was necessary for burst generation in L5B FS cells. Importantly, FS-FS coupling via either electrical or chemical synapses was unlikely to be disrupted by the cut, as FS cells in L5B do not, in general, extend dendrites into L4 (Gouwens et al., 2020), so this manipulation should not have affected oscillations driven by FS-FS interactions. It could be argued that the results of our manipulation are also consistent with the interpretation that L4→L5 connections were required only to provide tonic excitation to FS interneurons (Stark et al., 2014), rather than drive it on a cycle-to-cycle basis. However, FS cells in the cut slices still received strong excitation from the lower tier of thalamic terminations in L5, as evident by their undiminished first-spike response (Fig. 6B), so it is not clear why additional tonic excitation would have been needed. Even more importantly, our voltage-clamp data (Fig. 5A,C) revealed cycle-by-cycle, rhythmic EPSPs in RS interneurons occurring just before spikes in FS interneurons, suggesting that both cell types received EPSPs from the same presynaptic axons or at least from a highly synchronous presynaptic population. It is difficult to see how such precisely timed, rhythmic excitatory input would not be a driving factor in the entrainment of FS cells to the ripplet oscillation.

### The role of electrical or chemical FS-FS connections

FS cells in the rodent neocortex and hippocampus are known to be coupled by electrical synapses. These are prominent in juvenile rats, with coupling coefficients of 5-10% (Galarreta and Hestrin, 1999; Gibson et al., 1999, 2005; Otsuka and Kawaguchi, 2013). In the mouse, however, electrical coupling is much weaker, with reported coupling coefficients of 1.5% in juvenile (Hu et al., 2011; Hu and Agmon, 2015) and adult (Galarreta and Hestrin, 1999, 2002; Meyer et al., 2002) mice, although one study in juvenile mice (Deans et al., 2001) found a higher strength of coupling. In our current dataset of subadult and adult animals, the coupling coefficient between paired L4 FS cells, when detectable, was <1%. Interestingly, a recent ultrastructural study of gap junctions in barrel cortex has identified a subset of PV interneurons’ (“Type 1”), with somata located more centrally in the barrel and dendritic arbors fully contained within the barrel, which do not make gap junctions with each other but do make them with PV cells closer to the barrel periphery (Shigematsu et al., 2019). It is possible that our paired recordings, which were biased towards neighboring FS cells, preferentially sampled unconnected Type 1 pairs. Even so, the pairs we recorded were exquisitely synchronized. Our data therefore suggest that electrical coupling did not play a major role in the submillisecond FS-FS synchrony we observed, and by extension in the generation of ripplets. The conclusion that electrical coupling between FS interneurons was not a major factor in FS synchrony and ripplet generation is consistent with previous studies of hippocampal ripples, in which FS-FS electrical coupling is abolished but hippocampal ripples are largely unaffected in Connexin 36 (Cx36) knockout mice (Hormuzdi et al., 2001; Buhl et al., 2003). Notwithstanding, we cannot rule out that dendro-dendritic gap junctions, which have been described between PV-expressing interneurons (Katsumaru et al., 1988; Fukuda and Kosaka, 2000, 2003; Fukuda, 2017) and which may not be evident in somatic recordings, played a supporting role in synchronizing FS spiking during ripplets, in conjunction with GABAergic synapses (Gibson et al., 1999; Tamas et al., 2000; Szabadics et al., 2001; Hu and Agmon, 2015) or with shared, synchronous EPSPs (Galarreta and Hestrin, 2001; Hjorth et al., 2009).

In contrast to electrical coupling, FS-FS cells in the rodent cortex are often connected chemically, by one-way or reciprocal GABAergic synapses (Gibson et al., 1999; Deans et al., 2001; Galarreta and Hestrin, 2002; Ma et al., 2012; Hioki et al., 2013), and such chemical coupling can synchronize neurons with sub-millisecond precision (Hu et al., 2011). In our current dataset only about half of the recorded FS-FS pairs were connected chemically, whereas all pairs exhibited sub-millisecond firing synchrony during ripplets, arguing against a requirement for direct inhibitory synapses in this synchrony. By elimination, therefore, our data point to shared excitatory inputs as the main generator of submillisecond FS-FS synchrony during ripplets. While the initial common input originated in the thalamus, synchronization of subsequent spikes in the burst must have been driven by local, recurrent excitation. Similar conclusions were reached from observations of sharp synchrony between FS interneurons in the awake rabbit (Swadlow et al., 1998) and during cortical UP states in mice (Neske and Connors, 2016).

### Ideas and Speculations: Functional significance of ripplets

While our experiments were conducted ex-vivo, similar earlier reports (reviewed above) of ripplet-like LFP oscillations in vivo, in four other species (including humans) and in two cortical areas, suggest that ripplets may be a ubiquitous cortical response to a brief but salient sensory stimulus. In our experiments, the light pulse likely activated most, if not all thalamocortical terminals impinging on the recorded neurons. Such a massively synchronous input is not necessarily non-physiological: thalamocortical neurons display strong synchrony when activated by a sensory stimulus that also elicits a strong cortical response (Alonso et al., 1996; Pinto et al., 2000; Roy and Alloway, 2001; Temereanca and Simons, 2003; Bruno and Sakmann, 2006; Wang et al., 2010; Bale et al., 2015; Whitmire et al., 2016; Wright et al., 2021). Moreover, based on electrophysiological evidence, a high divergence/convergence of thalamocortical axon onto L4 FS interneurons has been proposed, approaching a “complete transmission line” with all-to-all connectivity (Swadlow, 1995; Bruno et al., 2003; Bereshpolova et al., 2020). Thus, our optogenetic activation was a reasonable approximation to a salient sensory stimulus, one which would activate a high fraction of L4 FS interneurons and which would be readily perceived by the animal. A testable prediction is therefore that in the behaving animal, stimuli which elicit ripplets will more likely be perceived by the animal, compared with stimuli which do not.

It is possible that ripplets are a stereotyped response to any strong sensory input, and that the spatiotemporal pattern of firing during a ripplet is not stimulus-specific. Alternatively, ripplets could provide the cortex with increased bandwidth for transmission and storage of specific sensory information. Similar to the encoding scheme proposed for gamma/theta oscillations (Lisman and Buzsaki, 2008; Lisman and Jensen, 2013) or for odor coding in the insect olfactory system (Laurent and Davidowitz, 1994), ripplets may provide a framework for clustering excitatory neuron spikes into discrete “time bins”, with each ripplet cycle providing a time bin of ~2.5 ms. While FS interneurons fire in every bin, excitatory neuron fire more sparsely, thus generating a binary sequence of 1’s (spike) and 0’s (no spike). Moreover, different excitatory neurons may generate different sequences. These binary sequences will be precisely repeated every time the same sensory stimulus is encountered, but may vary between different stimuli in tune with the temporal (Arabzadeh et al., 2005) or spatial (Andermann et al., 2004) variations elicited in the activity pattern of primary sensory, brainstem and thalamocortical neurons along the chain of activation from periphery to cortex. The assignment of a precise spike sequence to a neuron, rather than merely a total spike count, more than doubles the amount of information that can be conveyed by the neuron (Arabzadeh et al., 2006). Interestingly, three different studies examining the precision of the temporal code for stimulus location in rat somatosensory cortex (Ghazanfar et al., 2000; Panzeri et al., 2001; Foffani et al., 2004) found that peak discrimination, or maximal information content, is achieved when spikes are aggregated into time bins of 2-3 ms, which is exactly the period of a ripplet.

How will the information encoded in these putative binary patterns be read-out by downstream cortical targets? L4 excitatory cells make synapses on basal dendrites of L2/3 pyramidal neurons (Feldmeyer et al., 2002; Lubke et al., 2003; Shepherd and Svoboda, 2005), and L2/3 neurons are tuned to precise spatiotemporal sequences of synapse activation (Branco et al., 2010; Branco and Hausser, 2011). One can therefore envision that each spatiotemporal spike pattern in L4 cells will specifically activate those L2/3 pyramidal cells which are tuned to this exact spatiotemporal pattern of synaptic inputs. In this manner, ripplets can transform a temporal code representing a sensory feature, e.g. a texture, into a spatial code, with different ensembles of L2/3 pyramidal cells encoding different textures. Such ensembles can stabilize by Hebbian or spike time-dependent plasticity mechanisms and form “engrams”, long-term assemblies which can later be activated to recall the same sensory events which initially shaped them (Liu et al., 2012; Carrillo-Reid et al., 2019). That ripplets participate in the encoding of sensory information into transient or stable ensembles is a testable hypothesis.

### Summary and Future Directions

Our experiments revealed the synaptic interactions underlying ripplets - transient, ultrafast network oscillations evoked in the somatosensory cortex by a brief, synchronous thalamocortical volley. Previously observed in human EEG/MEG recordings and in a handful of animal intracellular studies, ripplet-like oscillations in sensory cortices may be much more frequent and widespread than previously recognized. The robustness and precise reproducibility of optogenetically evoked ripplets, and the accessibility of our ex-vivo experimental model to recording and imaging and to pharmacological and optogenetic manipulations, will allow future studies to examine the cellular and synaptic mechanisms of ripplets in exquisite detail and to construct, test and refine detailed computational models of the underlying L4 network which can faithfully simulate ripplet oscillations. In addition, studies of neocortical ripplets are likely to shed new light on the mechanisms behind similar oscillations in other brain areas, such as hippocampal ripples. Beyond mechanisms, the role of ripplets in encoding and transmitting sensory information is a highly intriguing topic awaiting future investigations.

## MATERIALS AND METHODS

### Animals

Animals used in this study were housed at the AAALAC-accredited WVU Lab Animal Research Facility according to institutional, federal and AAALAC guidelines. Animal euthanasia for brain slice preparation and for perfusion fixation was carried out under deep anesthesia. Animal use followed the Public Health Service Policy on Humane Care and Use of Laboratory Animals, and was approved by the WVU Institutional Animal Care and Use Committee (protocol #1604002316). West Virginia University has a PHS-approved Animal Welfare Assurance D16-00362 (A3597-01).

We crossed KN282 mice (MMRRC strain 036680-UCD), in which neurons in the ventrobasal (VB) complex of the thalamus express Cre recombinase (Gerfen et al., 2013; Hu and Agmon, 2016), with the Ai32 channelrhodopsin (ChR2) reporter (strain #024109, The Jackson Laboratory, Bar Harbor, ME, USA) (Madisen et al., 2012) or with the Ai9 tdTomato reporter (strain #007909) (Madisen et al., 2010). Dual-transgenic progeny of both sexes, 3-7 weeks old, were used for experiments. All mouse lines used were maintained for multiple generations as heterozygotes, by breeding mutant males with outbred (CD-1, Charles River Laboratories, USA) wild-type females.

### Slice preparation

Mice were decapitated under deep isoflurane anesthesia, the brains removed and submerged in ice-cold, sucrose-based artificial cerebrospinal fluid (ACSF) containing the following (in mM): Sucrose 206, NaH_2_PO_4_ 1.25, MgCl_2_.6H_2_O 10, CaCl_2_ 0.25, KCl 2.5, NaHCO_3_ 26 and D-glucose 11, pH 7.4. Thalamocortical brain slices (Agmon and Connors, 1991; Porter et al., 2001) of somatosensory (barrel) cortex, 300-350 μm thick, were cut in same solution using a Leica VT-200 vibratome, and placed in a submersion holding chamber filled with recirculated and oxygenated ACSF (in mM: NaCl 126, KCl 3, NaH_2_PO_4_ 1.25, CaCl_2_ 2, MgSO_4_ 1.3, NaHCO_3_ 26, and D-glucose 20). Slices were incubated for at least 30 minutes at 32°C and then at room temperature until use. For recording, individual slices were transferred to a submersion recording chamber and continuously perfused with 32°C oxygenated ACSF at a rate of 2–3 ml/min.

### Electrophysiological recordings

Recording were done on an upright microscope (FN-1, Nikon) under a 40X water immersion objective. For whole-cell recordings, glass micropipettes (typically 5–8 MΩ in resistance) were filled with an intracellular solution containing (in mM): K-gluconate 134, KCl 3.5, CaCl2 0.1, HEPES 10, EGTA 1.1, Mg-ATP 4, phosphocreatine-Tris 10, and 2 mg/ml biocytin, adjusted to pH 7.25 and 290 mOsm. Layer 4 FS and RS cells were identified visually by soma size and shape and targeted for single or dual whole-cell recordings using a MultiClamp 700B amplifier (Molecular Devices, San-Jose, CA, USA). Upon break-in, cells were routinely tested by a standardized family of incrementing 600 ms-long intracellular current steps in both negative and positive directions. Upon strong depolarization, FS cells invariably reached steady-state firing rates of 150-300 Hz and their spike trains showed no or little frequency adaptation, while RS cells typically fired an initial fast doublet, followed by a slower, steady-state firing at <50 Hz (Fig. 2 – Figure Supplement 1). In post-hoc analysis, the same records were used in to extract multiple electrophysiological parameters for each cell (Fig. 2 – Source Data 1). In dual-recording experiments, synaptic connectivity was routinely tested in both directions by stimulating one cell with a 10 Hz train of brief suprathreshold current steps and recording the response in the paired cell (at resting potential if the putative synapse was excitatory, or at −50 mV if inhibitory). Extracellular local field potentials (LFPs) were recorded using glass micropipettes filled with 0.9% sodium chloride, with a tip broken under visual control (tip diameter ~5 μm, 2-3 MΩ resistance), positioned in the center of the L4 barrel. The extracellular signal was amplified 1000× and low-pass filtered at 1.3 KHz using a differential amplifier (EXT-02F, NPI, Tamm, Germany); 20-100 trials were averaged. To block fast excitatory synaptic transmission, 6-cyano-7-nitroquinoxaline-2,3-dione disodium (CNQX, Tocris, Minneapolis, MN, USA) and d-(−)-2-amino-5-phosphonopentanoic acid (APV, Tocris) were added to the ACSF, at final concentrations of 20 and 30 μM, respectively. Data were acquired at a 20 KHz sampling rate using a National Instruments (Austin, TX, USA) ADC board and in-house acquisition software written in the LabView (National Instruments) environment.

### Data inclusion/exclusion

Intracellular recordings not passing our minimal quality criteria (resting potential ≤−60 mV (FS cells) or ≤−65 mV (RS cells) with no holding current, action potential peak ≥10 mV (FS cells) or ≥20 mV (RS cells)), were excluded from further analysis and reporting. Inclusion criteria for each analysis are stated in the Results section. Otherwise, no datapoints (spikes, cells, cell pairs or slices) within the stated inclusion criteria were excluded from the analysis, figures or reported results.

### Optogenetic stimulation

In most experiments, to activate thalamocortical axons and terminals, pulses of white light from a high-power LED light source (Prizmatix, Holon, Israel) were delivered through the epi-illumination light path and a GFP filter cube. The dual-color experiments related to Fig. 6 were done using an X-Cite multi-LED light source (Excelitas, Waltham, MA, USA). Maximal illumination intensity of the Prizmatix light source was 1.0 mW, as measured in air at the focal plane of the 40X objective. Assuming the full light energy was spread evenly over the field of view of the 40X objective (550 μm diameter) and no light scattering in water, calculated maximal illuminance at the tissue was 4.2 mW/mm^2^. In the course of the experiment, light intensity was adjusted to 10-90% of maximal intensity. Stimuli at each light level were typically repeated 8-12 times at 6–10 s intervals.

### Synchrony analysis

Given any two spike trains, synchrony is defined as the fraction of spikes in the shorter train occurring within a pre-determined “synchrony window” (SW) before or after a spike in the longer train. Chance synchrony is the expected (i.e. average) synchrony remaining after shifting each spike in the shorter train by a random jitter ≤J, J being the “jitter window”, and is calculated analytically rather than by Monte-Carlo simulations. The JBSI is the difference between the observed and chance synchrony, i.e. the magnitude of synchrony destroyed by the jitter, multiplied by a normalization factor β which limits the JBSI to values between 1 (highest possible synchrony) to −1 (lowest possible synchrony), with 0 indicating purely chance synchrony. Here we used J=2·SW and β=2. Unlike many other synchrony measures, the JBSI is independent of firing rates and firing rate differentials, and is not sensitive to slow co-modulations in these rates. See (Agmon, 2012) for theoretical and computational details. JBSI computations were done using in-house MathCad routines (MathCad, PTC). The routines, as well as a MatLab function, are available under the MIT license on https://github.com/aricagmon/JBSI-codes.

### Statistical analysis

Unless noted otherwise, exact p-values were computed using distribution-free, non-parametric permutation tests (Good, 1999) by performing 10,000 random permutations of the data and calculating the fraction of permutations resulting in equal or more extreme values of the relevant statistic (in both tails). When no more extreme values were found, this is indicated as p<0.0001. All data are reported as mean±SEM unless indicated otherwise.

## ACKNOWLEDGEMENTS

We are indebted to Qingyan Wang for outstanding technical support. We thank Barry Connors and Liset Menendez de la Prida for critical reading of the manuscript and helpful discussions. Craig Atencio translated the JBSI algorithm into MatLab code.

## FUNDING

This study was supported by National Institutes of Health grant NS116604 to AA. Additional funding was provided by Transition Grant Support from the Office of Research and Graduate Education, WVU Health Sciences Center, and by the Program to Stimulate Competitive Research, Office of Research, WVU. REH was supported by NIH training grants GM081741 and GM132494. Confocal imaging was performed at the WVU Microscopic Imaging Facility which was supported by NIH grants GM121322, GM103434, GM103503, GM104942, and RR016440.

## COMPETING INTERESTS

The authors declare no competing interests.

## Figure Supplements

Fig. 1- Figure supplement 1

Fig. 1- source data 1: *Fig1-SourceData1.xlsx*

Fig. 2 – Figure supplement 1

Fig. 2 – source data 2: *Fig2-SourceData1.xlsx*

**Fig. 1 – Figure supplement 1:**
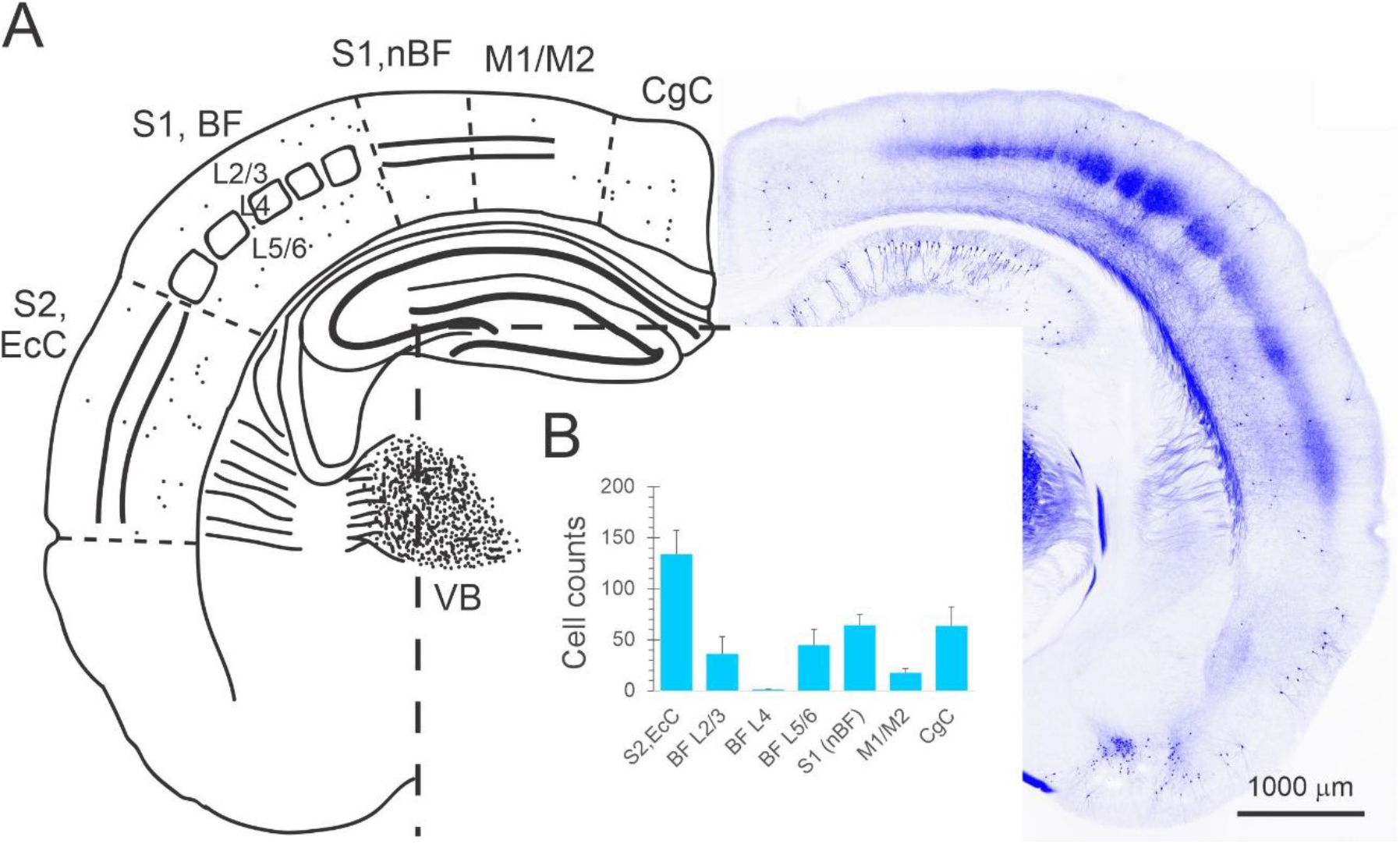
*C*re-expressing neurons in the KN282 mouse. **A, right**, a confocal projection of a representative 60 μm section through a KN282 x Ai9 brain, in which cre-expressing neurons were detected by their tdTomato fluorescence. **Left**, an outline sketch of the same section with counted cells indicated as dots, and with the different areas and layers counted indicated. CgC, cingulate cortex. M1/M2, primary/secondary motor cortex (also including parietal association areas in some anteroposterior levels). S1, nBF, primary somatosensory cortex, non-barrel field areas (e.g. forelimb and trunk areas). BF, barrel field. S2, EcC, secondary somatosensory cortex and ectorhinal cortex (also including temporal association areas and secondary auditory cortex in some anteroposterior levels). VB, ventrobasal thalamus. **B**, average cell counts from 4 animals, summed over five 60 μm sections per animal, sampled at 240 μm intervals through the barrel cortex (see Methods). Error bars represent sample standard deviation.

### Methods

Four male mice (4-6 week old) were deeply anesthetized with 2.5% Avertin and transcardially perfused with saline followed by 50 ml 4% paraformaldehyde (PFA). Brains were removed, post-fixed in 4% PFA at room temperature for 4 hr, stored for at least 72 hr in 30% sucrose solution in phosphate buffer saline (PBS) at 4° C, and sectioned coronally on a freezing stage sliding microtome. For imaging, five 60 μm sections were sampled per brain at 240 μm intervals, in total extending 800-2000 μm posterior to bregma, as determined by comparison with the Paxinos and Franklin atlas (Paxinos and Franklin, The Mouse Brain in Stereotaxic Coordinates, 4^th^ edition, Academic Press 2012). Confocal image stacks were taken with a 10× objective at 2.5 μm Z-steps on a Zeiss LSM 710 or a Nikon A1R confocal microscope. Labeled cells were counted by visual inspection of the full stack and summed per area over the 5 sections from each animal; counts thus represent the number of cells in a composite 300 μm section. Areal boundaries follow the designations in the Paxinos and Franklin atlas.

**Fig. 1 - Source data 1**: *Fig1_SourceData1.xlsx*

**Fig. 2 – Figure supplement 1:**
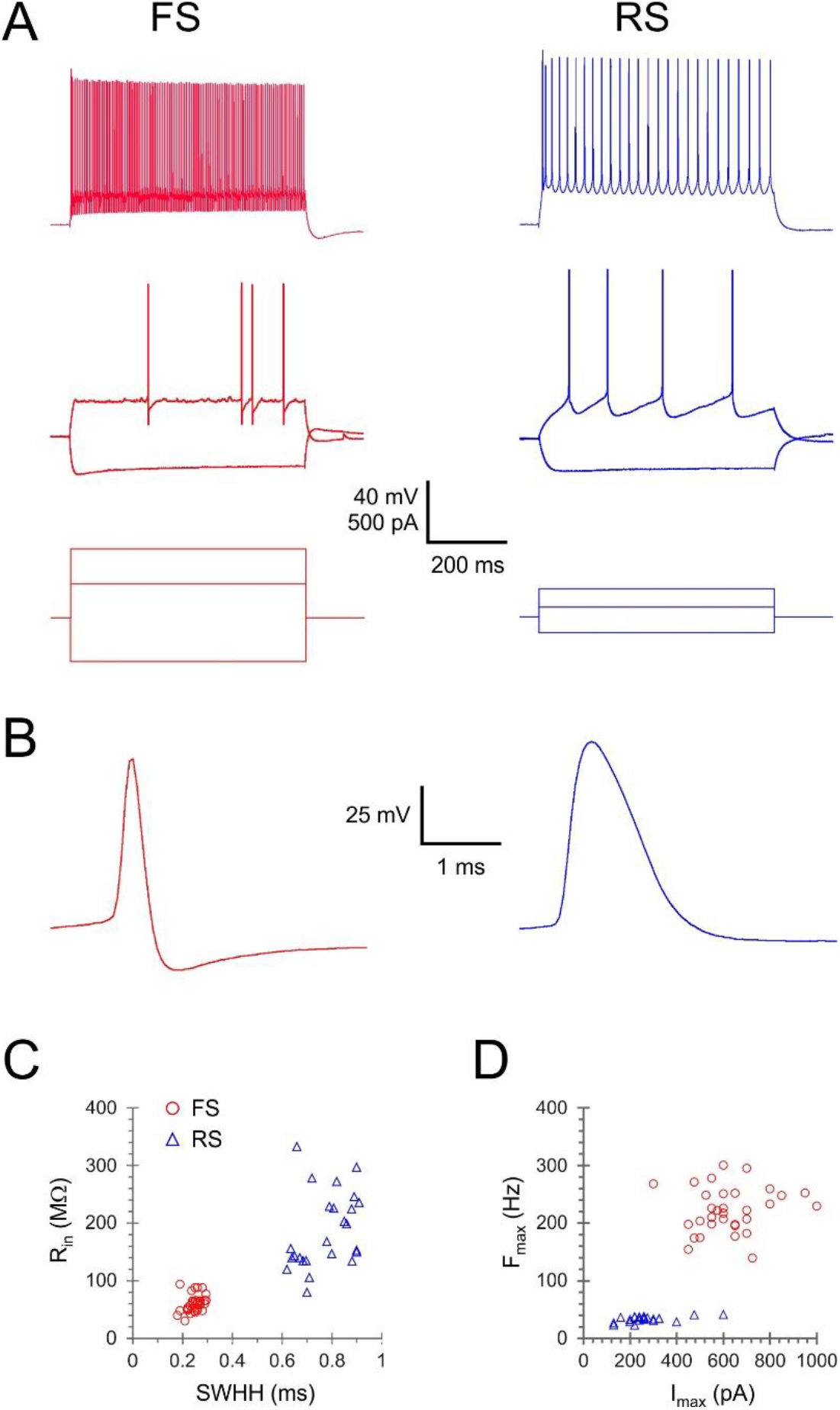
Comparison of electrophysiological parameters of FS and RS neurons. **A**, sub-and suprathreshold intracellular voltage responses (top and middle panels) to 3 levels of intracellular current steps (lower panel) in example FS and RS neurons. Note that FS cells achieved much higher steady-state firing frequencies, and that they required higher current steps compared with RS neurons reflecting their lower input resistance. **B**, single spikes plotted at an expanded timescale, revealing the pronounced difference is spike width and shape. **C**, input resistance (R_in_) plotted against spike width at half-height (SWHH) in the analyzed subset of animals (see Methods). **D**, maximal firing frequency (F_max_) plotted against the current step (I_max_) that was used to elicit it, in the same neurons as C.

#### Methods

Analysis was done on a subset of 32 FS interneurons and 25 RS cells in slices from 25 and 22 animals, respectively, 3-6.5 weeks old (14 animals were common to both subsets). A total of 8 electrophysiological parameters were analyzed per cell (summarized in Fig. 2 - Source Data 1).

#### Electrophysiological parameters definition

Single-spike parameters were measured at rheobase (minimal current evoking an action potential). All current steps were 600 ms long.

**V_rest_:** Resting potential upon break-in, with no holding current applied.

**V_threshold_:** The voltage where dv/dt=5 V/s.

**Spike height:** Spike peak - V_threshold_.

**Spike width at half-height (SWHH)**: spike width measured half-way between Vthreshold and spike peak.

**AHP**: V_threshold_ - Spike trough.

**R_in_**: The slope of the I-V plot, calculated from 4-6 positive and negative subthreshold current steps, at membrane potentials up to ±15 mV from rest.

**F_max_**: The steady-state firing frequency, computed as the reciprocal of the average of the last 5 ISI’s in a spike train elicited by I_max_.

**I_max_**: The maximal current step applied before a noticeable reduction in spike height.

**Fig. 2 - Source data 1**: *Fig2_SourceData1.xlsx*

## Notes

### Competing Interest Statement

The authors have declared no competing interest.

